# A tale of two tumors: differential, but detrimental, effects of glioblastoma extracellular vesicles (EVs) on normal human brain cells

**DOI:** 10.1101/2024.04.08.588622

**Authors:** Mary Wang, Arin N. Graner, Bryne Knowles, Charlotte McRae, Anthony Fringuello, Petr Paucek, Michael Gavrilovic, McKenna Redwine, Caleb Hanson, Christina Coughlan, Brooke Metzger, Vince Bolus, Timothy Kopper, Marie Smith, Wenbo Zhou, Morgan Lenz, Aviva Abosch, Steven Ojemann, Kevin O. Lillehei, Xiaoli Yu, Michael W. Graner

## Abstract

Glioblastomas (GBMs) are dreadful brain tumors with abysmal survival outcomes. GBM EVs dramatically affect normal brain cells (largely astrocytes) constituting the tumor microenvironment (TME). EVs from different patient-derived GBM spheroids induced differential transcriptomic, secretomic, and proteomic effects on cultured astrocytes/brain tissue slices as GBM EV recipients. The net outcome of brain cell differential changes nonetheless converges on increased tumorigenicity. GBM spheroids and brain slices were derived from neurosurgical patient tissues following informed consent. Astrocytes were commercially obtained. EVs were isolated from conditioned culture media by ultrafiltration, ultraconcentration, and ultracentrifugation. EVs were characterized by nanoparticle tracking analysis, electron microscopy, biochemical markers, and proteomics. Astrocytes/brain tissues were treated with GBM EVs before downstream analyses. EVs from different GBMs induced brain cells to alter secretomes with pro-inflammatory or TME-modifying (proteolytic) effects. Astrocyte responses ranged from anti-viral gene/protein expression and cytokine release to altered extracellular signal-regulated protein kinase (ERK1/2) signaling pathways, and conditioned media from EV-treated cells increased GBM cell proliferation. Thus, astrocytes/brain slices treated with different GBM EVs underwent non-identical changes in various ‘omics readouts and other assays, indicating “personalized” tumor-specific GBM EV effects on the TME. This raises concern regarding reliance on “model” systems as a sole basis for translational direction. Nonetheless, net downstream impacts from differential cellular and TME effects still led to increased tumorigenic capacities for the different GBMs.

## 1. Introduction

Glioblastomas (aka, CNS5 World Health Organization (WHO) Grade 4 isocitrate dehydrogenase (IDH)-wild type astrocytoma; we will abbreviate as GBMs), are the most common and deadliest of malignant brain tumors in adults with overall survival of 15-16 months, and near-universally fatal outcomes (Lukas *et al*., 2019; Ostrom *et al*., 2021; Wirsching *et al*., 2016). Despite the cataloging of genetic alterations, expanded target identities, new treatment modalities, and in particular, continued immunotherapeutic approaches (Cruz *et al*., 2022; Farrell *et al*., 2020; Yu and Quail, 2021), we have barely budged the needle for improved survival since the introduction of temozolomide with radiation as the standard of care in 2005 (Poon *et al*., 2020; Stupp *et al*., 2005),. Further, the disease inflicts physical, cognitive, and economic damage to patients and families during the ordeal (Gabel *et al*., 2019; Goel *et al*., 2021; Raizer *et al*., 2015). A better understanding of the tumor biology may enable better therapeutic intervention. Here, we focus on the diverse effects of GBM extracellular vesicles (EVs) as manipulators of the tumor microenvironment (TME) where the scenarios ultimately benefit the tumor.

Extracellular vesicles/nanoparticles are released from all cell types; these materials range from membrane-less “supermeres” (<30nm) and “exomeres” (30-50nm) (Anand *et al*., 2021; Jeppesen *et al*., 2022; Tosar *et al*., 2022; Zhang *et al*., 2018; Zhang *et al*., 2019a; Zhang *et al*., 2021) to membrane-enclosed vesicles such as “small EVs” (50-150nm) and “medium” or “large” EVs (>200nm to 1000nm) (Skovronova *et al*., 2021; Yanez-Mo *et al*., 2015). These size ranges, like the historical EV nomenclature related to biogenesis (“exosomes”, “microvesicles”), remain relatively ambiguous (Graner, 2019). We will use “EVs” throughout the rest of this work to refer to our isolated extracellular materials.

Tumor EVs have been long known to modulate the local TME (Clancy and D’Souza-Schorey, 2023), and GBM EVs play similar roles in that unique environment (Broekman *et al*., 2018; Graner, 2019; Matarredona and Pastor, 2019; Mondal *et al*., 2017). For high-grade gliomas such as GBMs, the bulk of cells constituting the normal – and tumor – microenvironment are astrocytes (Brandao *et al*., 2019; Graner, 2019; Nieland *et al*., 2021). We (Oushy *et al*., 2018) and others (Gao *et al*., 2020; Taheri *et al*., 2018; Yu *et al*., 2018; Zeng *et al*., 2020) have demonstrated impacts of GBM EVs on astrocytes (and other brain cells), and there are indications that astrocyte responses as recipients of GBM EVs end up generating a tumor-promoting environment (Hallal *et al*., 2019; Oushy *et al*., 2018). In this work we extend those studies to include a deeper molecular analysis of the astrocytes that were treated with EVs from two different GBM spheroid lines, along with functional read-outs. We show that the EVs from the different GBMs induce quite different changes in the recipient astrocytes leading to differential downstream effects, although there are similarities in some cases. We show this through transcriptomic and proteomic studies, secretome studies, proliferation assays, phospho-signaling arrays, and proteolysis assays. We find that EVs from the F3-8 GBM cells induce a viral-like response in astrocytes, while EVs from G17-1 GBM cells show more effects in the astrocyte extracellular space. Our work highlights the individual nature of GBM impacts on cells of the TME, and poses a cautionary frame for development of single-tumor model systems. Our work also suggests that true precision medicine may require extensive analyses of tumors, their EVs, and their influences on the different cell types as constituents of the TME.

## 2. Materials and Methods

### 2.1. Human subjects

The study has been approved by the Colorado Multiple Institutional Review Board (COMIRB, #13–3007). Tumor samples were collected from patients consented to the protocol prior to resection for brain tumors or epilepsy surgeries in the University of Colorado Hospital Department of Neurosurgery. The tissue samples were processed and stored at the Nervous System Biorepository at the Department of Neurosurgery, University of Colorado Anschutz Medical Campus. Cell lines were grown out from these tumor tissues as previously described (Hellwinkel *et al*., 2015; Oushy *et al*., 2018). Normal brain tissues were obtained from epilepsy lesion resections where adjacent normal tissue was determined by Neuropathology.

### 2.2. Cell lines

F3-8 has been previously described (Oushy *et al*., 2018); at the time of assessment by Neuropathology, it was described as a WHO Grade 4 glioblastoma, atypical small cell variant, *IDH* wildtype (negative for *IDH1* mutation by immunohistochemistry). G17-1 was called a WHO Grade 4 glioblastoma, *IDH* wildtype (negative for *IDH1* mutation by immunohistochemistry). O(6)-methylguanine-DNA methyltransferase (*MGMT*) promoter methylation was indeterminant; it was negative for epidermal growth factor receptor (*EGFR*) amplification, but abnormal for trisomy and tetrasomy for chromosome 7p/7 centromere sequences. It was also borderline for loss of Phosphatase and tensin homolog (*PTEN*) and chromosome 10 centromere sequences. Both F3-8 and G17-1 were primary tumors.

Other lines used here included HEK-293 (human embryonic kidney cells), normal human epithelial cells, and cells grown from tumors collected for the Neurosurgery Nervous System Biorepository: M2-7, M6-7, and M16-8. M2-7 is an extremely unusual cell line that has its own manuscript in preparation. Briefly, it was a spinal metastasis (recurrent disease) of an adult embryonal rhabdomyosarcoma (eRMS). M6-7 is a cell line from a clinically recurrent Grade 4 astrocytoma, *IDH1* mutant (from immunohistochemistry). M16-8 is another unusual tumor cell line with its own manuscript in preparation. Briefly, it is a multiply recurrent anaplastic pleomorphic xanthoastrocytoma, WHO Grade 3. HEK-293 cells were from ATCC (CRL-1573; Manassas, VA, USA), and were cultured in EX-CELL® 293 serum-free medium (14571C; Sigma-Aldrich, St Louis, MO, USA). Human epithelial cell culture was described previously (Oushy *et al*., 2018)

Astrocytes. Normal human astrocytes were obtained from Lonza (24169, Cat # CC2565, lot # 0000289765, Lonza Inc., Anaheim, CA, USA) from ScienCell (Cat # 1800, lot #’s 30417 and 33464, ScienCell Research Laboratories, 1610 Faraday Ave, Carlsbad, CA 92008) and from Gibco (K1884, Cat # 2019-07-01, Lot # 2103630, Gibco Life Technologies/Thermo Fisher, Waltham MA, USA). Astrocytes were cultured in astrocyte basal growth media (ABM™, CC-3187) with astrocyte growth medium supplements (AGM CC-3186). Astrocytes were cultured in T-75 or T-175 flasks (Thermo, 156499/159910 Nunclon Delta EasY flasks; ThermoFisher, Waltham, MA, USA) at 1.5-2.0x10^6^ and 3.5-5.0x10^6^ cells per flask, respectively.

Tumor cells were grown as spheroids as described previously (Hellwinkel *et al*., 2015; Oushy *et al*., 2018). Following collection from surgery, tissue was diced with scalpels and mechanically dissociated with the butt of a syringe plunger. Cells were either cultured directly or were filtered through a 100 µm cell strainer (431752; Corning Life Sciences, Tewksbury, MA, USA). Cells were cultured under stem cell-like conditions in Neurobasal A medium supplemented with 2 mM L-glutamine (GlutaMAX supplement, 35050061; ThermoFisher) 1X B27 (17504044) and N2 (17502048), and 5 ng/ml basic fibroblast growth factor (bFGF/FGF2; PHG0368) and epidermal growth factor (EGF; PHG0313) (all from ThermoFisher). This medium is referred to as “NBA”. Cells were cultured in T-75 or T-175 flasks (ThermoFisher).

Brain tissues (parietal lobe) were collected in cold HGPSA medium (Hibernate A, 1X GlutaMax (Sigma), PenStrep, and Amphotericin B (all the rest from ThermoFisher)) sliced into ∼1mm cubes with scalpels, washed in NBA, and incubated in NBA (1.5ml) in 12-well plates (ThermoFisher).

### 2.3. Extracellular vesicle isolation and characterization

EVs were isolated from conditioned media via differential centrifugation, ultrafiltration/concentration, and ultracentrifugation, as described (Hellwinkel *et al*., 2015; Oushy *et al*., 2018). This is diagrammed in Supplemental Figure 1. EVs were characterized using NanoSight Nanoparticle Tracking Analysis (NTA), Exo-Check arrays, ExoCET assays, and transmission electron microscopy (TEM).

#### 2.3.1. NanoSight nanoparticle tracking analysis

EVs were diluted (1:100 or 1:1000) in particle-free phosphate buffered saline (PBS). EV numbers and diameters were determined using a NanoSight NS300 (Malvern Panalytical Ltd, Malvern, UK) with these analytical settings: camera type: sCMOS, level 11 (NTA 3.0 levels); green laser; slide setting and gain: 600, 300; shutter: 15 ms; histogram lower limit: 0; upper limit: 8235; frame rate: 24.9825 fps; syringe pump speed: 25 arbitrary units; detection threshold: 7; max. jump mode: auto; max. jump distance: 15.0482; blur and minimum track length: auto; first frame: 0; total frames analyzed: 749; temp: 21.099-22.015°C; and viscosity: 1.05 cP.

#### 2.3.2. Exo-Check arrays

These are antibody-capture arrays for semi-quantitative detection of 8 putative markers of exosomes/EVs (EXORAY200B, System Biosciences/SBI, Palo Alto, CA, USA). These were performed according to manufacturer’s instructions.

#### 2.3.3. ExoCET assays

This is an enzymatic (acetylcholinesterase, AChE) assay (EXOCET, Systems Biosciences/SBI) ostensibly used to quantitate exosomes/EVs based on AChE content. It uses a standard curve generated from exosomes/EVs of known quantity (based on NanoSight) supplied with the kit. We followed the manufacturer’s instructions.

#### 2.3.4. Transmission electron microscopy

Negative staining of EV samples and transmission electron microscopy (TEM). The negative staining and EM observation of EV samples were performed at the Electron Microscopy Center of the University of Colorado School of Medicine. The EV specimen (5 µl) was dropped onto a 200–400 mesh carbon/formvar coated grid and allowed to adsorb to the formvar for one minute. The 2% uranyl acetate (10 µl) was placed on the grid for one minute. Samples were rinsed in PBS for one minute. The sample was observed under a transmission electron microscope (FEI Tecnai G2 Biotwin TEM, Thermo Fisher, Waltham, MA, USA) at 80 KV with a magnification of 30,000 to 120,000x. The images were taken by the AMT camera system.

### 2.4. Treatment of astrocytes with EVs

Astrocytes were cultured as described above. EVs were diluted directly to 500 μg/ml in ABM medium and filtered through a sterile 0.45μm syringe filter. This concentration was based on our previous work (Oushy *et al*., 2018) and references cited therein. ABM medium was removed, and astrocytes were washed with PBS; EVs in ABM were then incubated with astrocytes for 24hrs. In some cases, astrocytes were treated with 1μM of the ERK1/2 inhibitor SCH772984 (S7101; Selleckchem/Selleck Chemicals LLC, Houston, TX, USA) for 4hrs prior to treatment with EVs. Conditioned media were collected and centrifuged at 1500 x g, 10min, RT, to remove cells and debris, and the media were either utilized in assays or were aliquoted and frozen at -80°C for future use. Cells were rinsed 3 times with PBS, and were harvested differently depending on the assay. For proteomics, cells were scraped directly off the plates in residual PBS. For Western blotting, cells were scraped in 1.0-3.0ml “radio-immune precipitation buffer” (RIPA, R0278; Sigma-Aldrich) supplemented with protease inhibitors and phosphatase inhibitors (cOmplete™ Mini, ethylenediaminetetraacetic acid (EDTA)-free Protease Inhibitor Cocktail, 4693159001; PhosSTOP™, 4906837001; Roche via Sigma-Aldrich). For particular kits, lysis buffers included in the kits were used per manufacturers’ instructions. Proteins were quantified by BCA Assay (Pierce™ BCA Protein Assay Kit, 23225; via ThermoFisher). Serum albumin was used as the standard.

Brain slices were treated in 1.5ml NBA media (with or without EVs) as described above. Media was collected at the end of the incubation and handled as described above. All values were normalized by weight of the brain slices.

### 2.5. Transcriptomics

Total RNA was extracted from astrocytes using RNeasy Plus Micro Kits (74034; QIAGEN, Germantown, MD, USA). RNA quality/quantity was checked with a 2100 Bioanalyzer (Agilent Technologies, Santa Clara, CA, USA). RINs were 9.9 or higher. Samples were then handled by the University of Colorado Anschutz Genomics and Microarray Core for microarray analysis using a Human Clariom D chip (902922; Applied Biosystems via ThermoFisher). This array has over 540,000 transcripts on it including mRNAs, miRNAs, lncRNAs, and splice variants. RNAs were analyzed using Transcriptome Analysis Console (TAC) 4.0.1 (ThermoFisher).

### 2.6. Mass spectrometry-based proteomic analysis

For astrocytes: after cell scraping in residual PBS (following PBS washes) and centrifugation (17,000 x g, 20min, 4°C) to pellet cells, cells were flash-frozen and stored at -80°C until use. For GBM spheres: cells/spheres were harvested by centrifugation (1500 x g, 5min, RT) and were washed with ice-cold PBS 3 times via centrifugation. Cell pellets were finally collected (17,000 x g, 20min, 4°C), and were flash-frozen and stored at -80°C until use. EVs were isolated as described above, and were flash-frozen before storage at -80°C until use. Samples were handled by the Mass Spectrometry Proteomics Shared Resource Facility on the CU Anschutz Campus.

#### 2.6.1. Sample digestion

The samples were digested according to the FASP (Filter-Aided Sample Preparation) protocol using a 10 kDa molecular weight cutoff filter. In brief, the samples were mixed in the filter unit with 200 µl of 8 M urea, 0.1M ammonium bicarbonate (AB) pH 8.0, and centrifuged at 14,000 g for 15 min. The proteins were reduced with 10 mM DTT for 30 min at room temperature (RT), centrifuged, and alkylated with 55 mM iodoacetamide for 30 min at RT in the dark. Following centrifugation, samples were washed 3 times with urea solution, and 3 times with 50 mM AB. Protein digestion was carried out with sequencing grade modified Trypsin (Promega) at 1/50 protease/protein (wt/wt) at 37°C overnight. Peptides were recovered from the filter using 50mM AB. Samples were dried in Speed-Vac and desalted and concentrated on Thermo Scientific Pierce C18 Tip.

#### 2.6.2. Mass spectrometry analysis

Samples were analyzed on an Orbitrap Fusion mass spectrometer (ThermoFisher) coupled to an Easy-nLC 1200 system (ThermoFisher) through a nanoelectrospray ion source. Peptides were separated on a self-made C18 analytical column (100 µm internal diameter, x 20 cm length) packed with 2.7 µm Cortecs particles. After equilibration with 3 µl 5% acetonitrile 0.1% formic acid, the peptides were separated by a 120 min linear gradient from 6% to 38% acetonitrile with 0.1% formic acid at 400 nL/min. LC mobile phase solvents and sample dilutions used 0.1% formic acid in water (Buffer A) and 0.1% formic acid in 80% acetonitrile (Buffer B) (Optima™ LC/MS, ThermoFisher). Data acquisition was performed using the instrument supplied Xcalibur™ (version 4.1) software. Survey scans covering the mass range of 350–1800 were performed in the Orbitrap by scanning from m/z 300-1800 with a resolution of 120,000 (at m/z 200), an S-Lens RF Level of 30%, a maximum injection time of 50 milliseconds, and an automatic gain control (AGC) target value of 4e5. For MS2 scan triggering, monoisotopic precursor selection was enabled, charge state filtering was limited to 2–7, an intensity threshold of 2e4 was employed, and dynamic exclusion of previously selected masses was enabled for 45 seconds with a tolerance of 10 ppm. MS2 scans were acquired in the Orbitrap mode with a maximum injection time of 35 milli seconds, quadrupole isolation, an isolation window of 1.6 m/z, HCD collision energy of 30%, and an AGC target value of 5e4.

#### 2.6.3. Protein identification and databases

MS/MS spectra were extracted from raw data files and converted into mgf files using Proteome Discoverer Software (ver. 2.1.0.62). These mgf files were then independently searched against the human database using an in-house Mascot server (Version 2.6, Matrix Science). Mass tolerances were +/- 10 ppm for MS peaks, and +/- 25 ppm for MS/MS fragment ions. Trypsin specificity was used allowing for 1 missed cleavage. Met oxidation, protein N-terminal acetylation, and peptide N-terminal pyroglutamic acid formation was allowed as variable modifications. Scaffold (version 4.8, Proteome Software) was used to validate MS/MS-based peptide and protein identifications. Peptide identifications were accepted if they could be established at greater than 95.0% probability as specified by the Peptide Prophet algorithm. Protein identifications were accepted if they could be established at greater than 99.0% probability and contained at least two identified unique peptides.

The partial least squares-discriminant analysis (PLS-DA) and heatmaps were generated using the MetaboAnalyst 5.0 online platform (https://www.metaboanalyst.ca/MetaboAnalyst/ModuleView.xhtml) with sum and range scaling normalizations. Transcriptomic and proteomic data were further analyzed using QIAGEN Ingenuity Pathway Analysis (IPA; QIAGEN Inc., Hilden, Germany https://digitalinsights.qiagen.com/IPA). We also used FunRich Functional Enrichment Analysis (http://www.funrich.org/) and STRING protein-protein interaction networks (https://string-db.org/) for some analyses.

### 2.7. Interferon (IFN) release assay

Astrocyte conditioned media were assayed for interferons and related cytokines using VeriPlex Human Interferon 9-Plex ELISA (51500; PBL Assay Science, Piscataway, NJ, USA) following manufacturer’s instructions. Spot intensities were quantified using an Alpha Innotech FluorChem Q System with AlphaView software (ProteinSimple/Bio-Techne, 3001 Orchard Pkwy, San Jose, CA, USA). Nine dots + controls per well were quantified per triplicate samples after converting pixel intensity data using Image J.

### 2.8. Western blotting

EV and control-treated astrocytes were harvested as described above. Equal volumes of sample and 2X Laemmli sample buffer (Bio Rad, cat # 1610737; Bio Rad Laboratories, Hercules, CA, USA) plus 2-mercaptoethanol (50 µl per 950 µl sample buffer, cat # 1610710, Bio Rad) were mixed to generate reducing and denaturing conditions. Samples were electrophoresed on Criterion™ TGX™ (Tris-Glycine eXtended) precast gels (10% Midi gels, cat # 5671033, Bio Rad) with Tris/Glycine/SDS running buffer (Bio Rad, cat # 1610732). Gels were electrotransferred with an iBlot 2 Dry Blotting System (cat # IB21001, ThermoFisher) onto nitrocellulose transfer stacks (cat # IB23001, ThermoFisher). Blots were rinsed in MilliQ water and stained with Ponceau S (cat # P7170, Sigma-Aldrich) to verify transfer and equal loading. Blots were rinsed in Tris-buffered saline (TBS, cat #1706435, Bio Rad) with 0.1% (vol/vol) Tween® 20 (Polysorbate 20) (cat #1706531, Bio Rad; buffer is called TBST). Blots were blocked with 5% non-fat dry milk (NFDM, Nestle Carnation code #1706531; Nestle USA, 28901 NE Carnation Farm Rd Carnation, WA, USA) (wt/vol) in TBST for 1hr at room temperature on a rocking platform. Blots were rinsed 3 times (5 minutes per wash) with TBST at room temperature while rocking. Blots were then incubated overnight at 4°C (while rocking) with the following primary antibodies: anti-OAS1 rabbit polyclonal 14955-1-AP; anti-OAS2 rabbit polyclonal 19279-1-AP; anti-OAS3 rabbit polyclonal 21915-1-AP; anti-MDA5/IFIH1 rabbit polyclonal 21775-1-AP; anti-RIG-I/DDX58 rabbit polyclonal 25068-1-AP; anti-IFIT3 rabbit polyclonal 15201-1-AP; all from Proteintech, used at 1:500 dilutions in 5% NFDM/TBST (Proteintech Group, Inc, 5500 Pearl Street, Suite 400, Rosemont, IL 60018, USA); anti-GFAP rabbit monoclonal #12389 (D1F4Q) (1:1000 dilution in 5% NFDM/TBST); anti-GAPDH rabbit monoclonal clone #5174 (D16H11) (1:5000 in 5% NFDM/TBST), both from Cell Signaling, Danvers, Massachusetts, USA. Blots were washed in TBST as described above. Secondary antibody was a goat anti-rabbit IgG with HRP label, #7074 from Cell Signaling, used at 1:5000 dilution in 5% NFDM/TBST. It was incubated on the blots on a rocker for 1hr at room temperature. Following washings as described, blots were treated with SuperSignal™ West Femto Substrate developer (cat # 34096; ThermoFisher) and bands were visualized using an Alpha Innotech FluorChem Q System (ProteinSimple/Bio-Techne). Protein/gene abbreviations used here: OAS = 2’,5’-oligoadenylate synthetase; MDA5 = melanoma differentiation-associated protein 5, IFIH1 = interferon induced with helicase C domain 1; RIG-1 = retinoic acid-inducible gene I, DDX58 = DExD/H-box helicase 58; IFIT3 = interferon induced protein with tetratricopeptide repeats 3; GFAP = glial fibrillary acidic protein; GAPDH = glyceraldehyde-3-phosphate dehydrogenase.

### 2.9. Cytokine release/‘secretome’ assays

Following treatment of astrocytes with EVs, the conditioned media were assayed with Proteome Profiler Human XL Cytokine Arrays (ARY022B; R&D Systems, Minneapolis, MN, USA) performed as per manufacturer’s protocols. Arrays were developed with chemiluminescence (SuperSignal™ West Femto Substrate, 34096; ThermoFisher) and spot intensities quantified using an Alpha Innotech FluorChem Q System (ProteinSimple/Bio-Techne). Background was subtracted from each membrane, and average luminosity from duplicate spots was compared between treatment groups. Results were display as heatmaps using Microsoft Excel (Microsoft 365; Microsoft, Redmond, WA, USA).

### 2.10. GBM cell proliferation in astrocyte-conditioned medium

G17-1 GBM cells were transferred from NBA to ABM medium and allowed to adapt for 7 days. Cells were harvested and grown in 24-well plates at 5 × 10^4^ cells/well. Conditioned media from astrocytes that were treated with 500 µg/ml G17-1 or HEK EVs (24hr) or PBS control were used to replace the old media. Conditioned astrocyte medium was centrifuged (1500g, 15 min) to remove any cells before adding the medium to the GBM cells. After 24hr, G17-1 cell proliferation was measured by MTS assay (Cell Titer 96 Aqueous One, Promega, Madison, WI, USA).

### 2.11. Phospho-signaling arrays

Astrocyte lysates from treated and control cells were incubated on Creative Biolabs Human Phospho-Kinase Antibody Array (AbAr-0225-YC; SUITE 203, 17 Ramsey Road, Shirley, NY 11967, USA) following the manufacturer’s instructions. Arrays were developed, quantified, and displayed as described above.

### 2.12. Periostin ELISA

Astrocyte culture supernatants were collected as described above following treatments of cells. Periostin (POSTN/OSF2) was measured in by ELISA (ELH-POSTN; RayBiotech, Peachtree Corners, GA, USA) according to manufacturer’s instructions.

### 2.13. Protease arrays

Conditioned media was collected from EV-treated or untreated (control) astrocytes and handled as described above. We incubated 1.5ml of media on Protease Arrays (R&D Systems Proteome Profiler Human Protease Array Kit, Cat # ARY021B; R&D Systems Inc., 614 McKinley Place NE, Minneapolis, MN 55413, USA), and followed the manufacturer’s instructions. Arrays were developed, quantified, and displayed as described above.

### 2.14. Protease assays

Matrix metalloproteinase (MMP) activities (as a general enzyme class) were assayed using abcam MMP Activity Assay Kits (Cat # ab112146, Fluorescent [green]; abcam, 152 Grove Street, Suite 1100 Waltham, MA 02453, USA) following manufacturer’s instructions. Samples were conditioned media from treated or control astrocytes, or from EVs directly. Measurements were performed under kinetic conditions for 0-1hr on a plate reader (excitation/emission = 490/525nm; Biotek Synergy H4 hybrid reader, Winooski, VT, USA, software version Gen5 3.09). We also used an EnzChek™ Gelatinase/Collagenase Assay Kit (cat # E12055; ThermoFisher) for protease assessment of brain slice conditioned media and EVs following manufacturer’s instructions. This was also a kinetic assay from 0-14hrs with fluorescent readouts (excitation/emission = 490/525nm; Biotek Synergy H4 hybrid reader).

### 2.15. Two-photon excitation microscopy of brain tissue

For 2-photon excitation imaging we used a laser scanning confocal microscope LSM 780 (Zeiss LS; Jena, Germany) equipped a two-photon tunable (690nm-1040nm) infrared pulsed laser MaiTai (SpectraPhysics, Milpitas, CA). The setup is equipped with an environmental chamber (37 °C, 5% CO2, and humidity control) for live tissue imaging. C-Apochromat 40x/1.20 W Korr FCS M27 objective was used for imaging of brain tissue excited with the two-photon laser tuned at 800 nm. Two-photon microscopy max intensity projection was obtained from Z-stack to visualize structural images in deep tissue. 3D Intensity projection (XY image size x: 1024, y: 1024, Z: 33, 8-bit) was reconstructed from Z-stack sample (x: 353.90 um, y:353.90 um, z: 10.65 um, 33 slides) scan mode with pixel dwell time of 1.58 us, open pinhole, spectral emission filter set to detect the whole visible range. 2D maximum intensity projection image size is 1024 x 1024 resolution.

### 2.16. Data analysis

Experiments were repeated at least once, often multiple times. Data were statistically assessed using Microsoft Excel and/or GraphPad Prism software (GraphPad Prism 9.0, San Diego, CA, USA). Student’s t-test or one-way or two-way ANOVA analyses were performed depending on experimental circumstances. ANOVAs were followed by pairwise multiple comparisons with Tukey’s method to show the significant differences between treatments and groups. Data were presented as mean ± SEM (or SD); significance was set at 0.05. In some cases, internal algorithm statistics were applied.

## 3. Results

### 3.1. GBM cell lines and their EVs

GBM cell lines F3-8 and G17-1 were generated from surgically-resected patient tumors and were grown in a stem cell fashion, resulting in “neurosphere” formation (Hellwinkel *et al*., 2015; Oushy *et al*., 2018). Curiously, many G17-1 spheroids would project long extensions onto the tissue culture plastic with mild adhesions while the F3-8 line grows as large (∼200μm) non-adherent clusters (Supplemental Figure 1A). EVs were isolated from the conditioned media by differential centrifugation, ultrafiltration, ultraconcentration, and ultracentrifugation (schematic in Supplemental Figure 1B). Nanosight nanoparticle tracking analysis shows expected sizes for “small” EVs (main peaks of 102 and 132nm for F3-8 and G17-1 EVs, respectively, Supplemental Figure 1C), and transmission electron microscopy reveals double membrane vesicles in 100-200nm diameters for EVs from both cell types, consistent with the Nanosight data (Supplemental Figure 1D). ExoCheck arrays indicate the presence of standard EV markers (cluster of differentiation 63 (CD63), annexin V (ANXA5), tumor susceptibility gene 101 (TSG101), flotillin 1 (FLOT1), intercellular adhesion molecule (ICAM), programmed cell death 6-interacting protein (PDCD6IP)/ALG-2 interacting protein x (ALIX), CD81) for both F3-8 and G17-1 EVs, along with characterization of M2-7, M6-7, and M16-8 EVs (Supplemental Figure 1E). The ExoCET assay, an acetylcholinesterase activity assay, is shown for HEK, F3-8, and G17-1 EVs; we do not regard this as a quantitative assay, but one to demonstrate the retention of biologic enzymatic activity following EV isolation by our protocol (Supplemental Figure 1F). These data indicate that we have isolated EVs from the cell lines based on established criteria.

### 3.2. Astrocyte transcriptome changes following GBM EV incubation

As a minimalist system to represent the GBM TME, we treated cultured astrocytes with GBM EVs (500μg/ml, 24hrs – dose and time were chosen based on maximum changes in signaling events from our previous work (Oushy *et al*., 2018), collecting conditioned media supernatant and cells for downstream analyses. Astrocytes receiving GBM EVs underwent large-scale changes in mRNA expression at the transcriptome level as evidenced in the volcano plots in Supplemental Figure 2 (control treatment in these cases consisted of EVs from normal epithelial cells). Ingenuity Pathway Analysis (IPA) condensed summaries are shown in Supplemental Figures 3 and 4, again with interferon (IFN) signaling and pathogen response pathways prominent in F3-8 EV-treated astrocytes (Supplemental Figure 3). G17-1 EV-treated astrocytes displayed pathways of cellular activation involving cyto/chemokine release, cell activation and migration, and potential changes in signaling pathways involving ERK1/2 (Supplemental Figure 4). Data from the Clariom D arrays are in Supplemental Tables 1 and 2.

### 3.3. Differential astrocyte proteome changes upon GBM EV incubation

To extend the transcriptomic findings we treated astrocytes with GBM EVs (control = PBS) and performed mass spectrometry-based proteomics on the cells. There, we found significant changes in astrocyte protein expression following GBM EV incubation compared to control (PBS-treated) cells as seen in volcano plots (Figure 1A, B) and hierarchical clustering heat maps (Figure 1C, D), indicating that both GBM cell lines’ EVs induced notable changes. This is further borne out in Ingenuity Pathway Analysis (IPA) where we see divergence in the astrocyte response to F3-8 vs G17-1 EVs (Supplemental Figures 5 and 6). The astrocyte response to F3-8 EVs is characterized by cyto/chemokine pathways involved in antiviral responses (Supplemental Figure 5) while astrocyte responses to G17-1 EVs involve cellular junctional pathways and signaling surrounding cellular transformation and migration (Supplemental Figure 6). This information mirrors what was seen in the transcriptomic analyses. IPA-generated Canonical Pathways for the F3-8 EV-treated astrocyte proteome highlight the cyto/chemokine influences driven by signal transducer and activators of transcription (STAT) activation and IFN-related molecular upregulation (Figure 2A). The Network in Figure 2B shows the connectivity of upregulated proteins intrinsic in innate anti-viral responses such as DDX58/RIG-I and the 2’-5’-oligoadenylate synthetases (OAS1-3), along with the IFN-inducible genes/proteins replete in the Network in Figure 2C. Significant Canonical Pathways arising from the G17-1 EV-treated astrocyte proteome include multiple junctional signaling pathways (from epithelial adherens, Sertoli cell, and gap junctions) with involvement of numerous structural proteins along with activated coronavirus replication pathways largely due to changes in tubulin expression (Figure 2D). The G17-1 EV-driven Network in Figure 2E implies an activated (janus kinase) JAK/STAT/ERK signaling axis where the reduced expression of the phosphatase protein tyrosine phosphatase receptor type E (PTPRE) may allow the activation of kinases in that node. We point out that IPA Network Legends (node and path design shapes, edges and their descriptions) can be found in Supplemental Figure 7. Proteins identified in the mass-spectrometry determinations are found in Supplemental Table 3.

**Figure 1:**
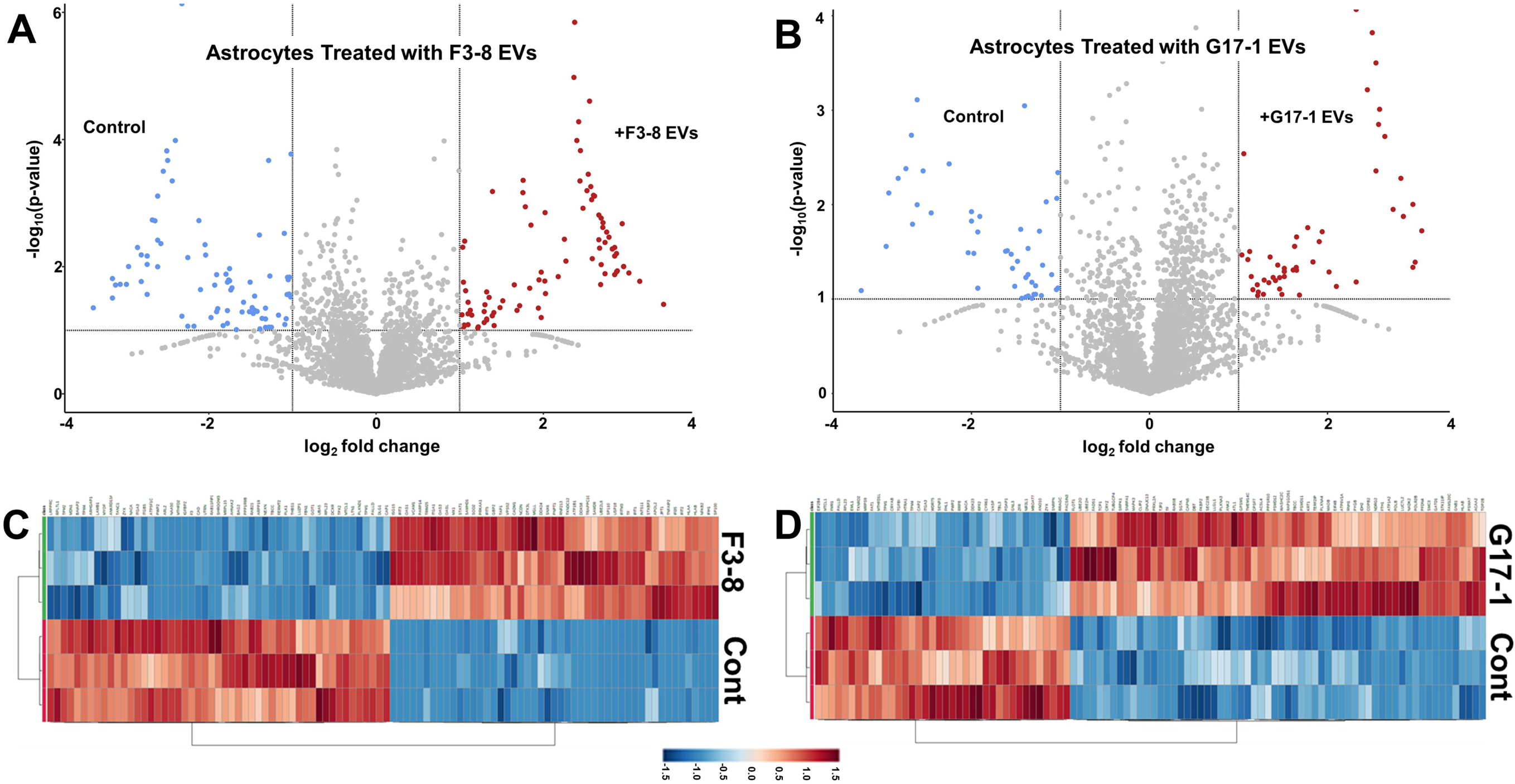
Proteomes of astrocytes treated with GBM F3-8 EVs or GBM G17-1 EVs. Data were generated in Metaboanalyzer 5.0. (A, B) volcano plots showing significantly differential proteomes between astrocytes treated with GBM EVs (upper right quadrants) or control treatments (PBS, upper left quadrants). (A) astrocytes treated with F3-8 EVs; (B) astrocytes treated with G17-EVs. (C, D) hierarchical clustering heatmaps from ANOVA statistical analyses utilized normalized data that were standardized by autoscaling features (top 100) with Euclidean distance measurements and clustered by ward. (C) Astrocytes treated with F3-8 EVs; (D) astrocytes treated with G17-1 EVs.

**Figure 2:**
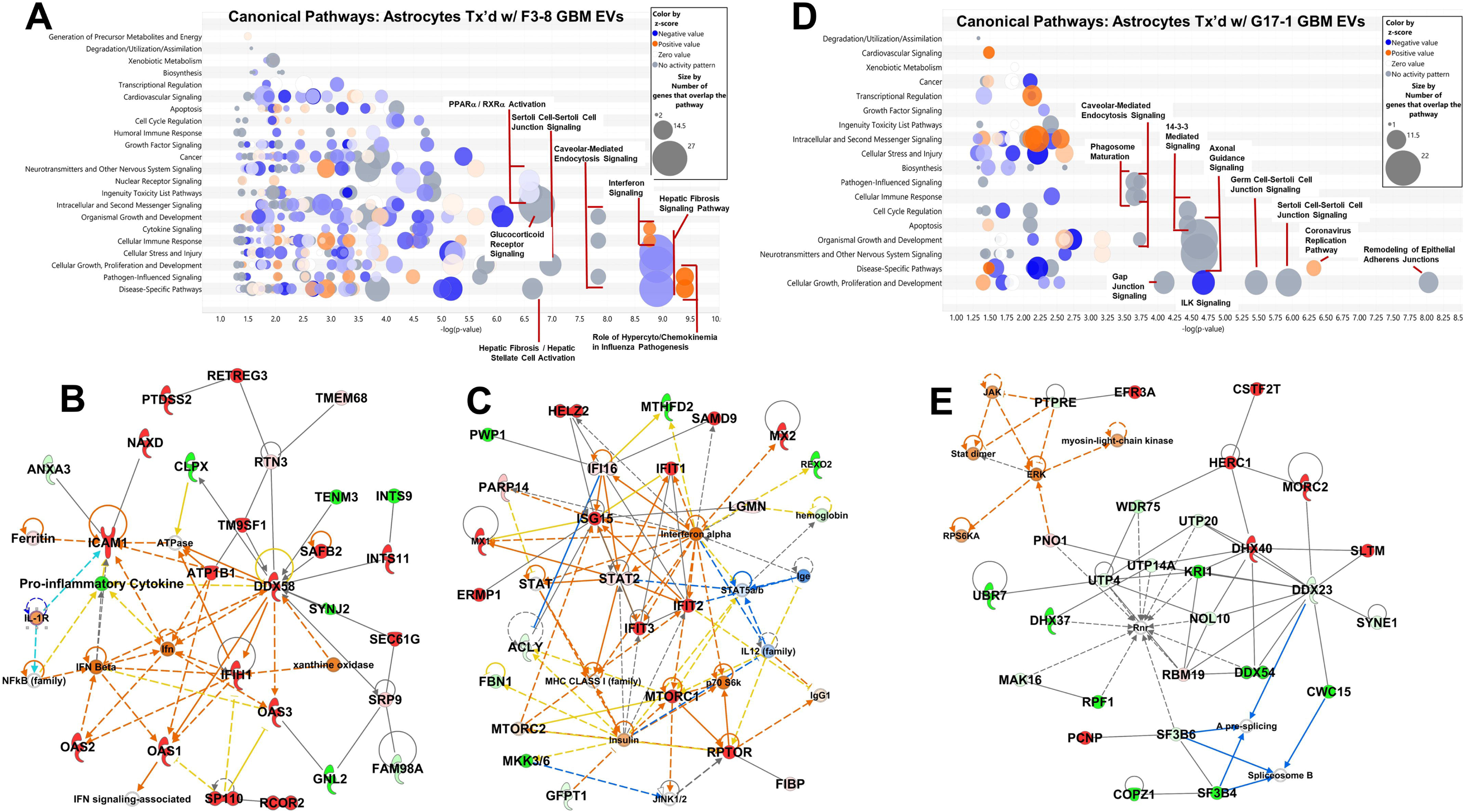
Ingenuity Pathway Analysis (IPA) highlights of astrocytes proteomes following treatment (“tx’d”) with GBM EVs: Canonical Pathways and relevant Networks. (A) “Bubble chart” of significant Canonical Pathways (based on -log(p-values)) derived from the proteome of astrocytes treated with GBM F3-8 EVs; broad categories are on the y-axis, and more specific categories are designated in the bubbles (higher scoring categories are denoted). Bubble sizes and colors scheme is in the inset. (B) Relevant interactome shown for F3-8 EV-treated astrocytes (Network 4 = Cardiovascular Disease; Infectious Disease; Organismal Injury and Abnormalities. Score = 44, 26 Focus Molecules). (C) Relevant interactome shown for F3-8 EV-treated astrocytes (Network 7 – Antimicrobial Response; Immunological Disease; Inflammatory Response. Score = 33, 21 Focus Molecules). (D) “Bubble chart” of significant Canonical Pathways derived from the proteome of astrocytes treated with GBM G17-1 EVs (as described in Fig 2A). (E) Relevant interactome shown for G17-1 EV-treated astrocytes (Network 4 - Connective Tissue Disorders; Developmental Disorder; RNA Post-Translational Modification. Score = 46, 27 Focus Molecules). “Scores” are based on Fisher’s exact test, -log(p-value); “Focus Molecules” are considered focal point generators within the network. Number of genes illustrated is limited to 35 by the algorithm. IPA Network Legends (node and path design shapes, edges and their descriptions) are in Supplemental Figure 7.

The transcriptomic and proteomic changes in GBM EV-treated astrocytes indicate that the different sources of GBM EVs lead to different astrocyte responses. We will dissect some of those distinct responses with functional assays focusing individually on astrocytes treated with either F3-8 or G17-1 GBM EVs.

### 3.4. Astrocytes innate anti-viral responses following incubation with GBM F3-8 EVs

Both the transcriptome and proteome profiles of astrocytes treated with F3-8 EVs strongly suggest responses to viruses/pathogens with upregulation of OAS1-3 genes and proteins, and upregulation of DDX58/RIG-I gene and protein (Figure 2B; IFIH1/MDA5 is also significantly upregulated but is not shown in that interactome). These are associated with sensing of intracellular RNAs and downstream signaling, driven by IFNs, or leading to IFN expression (Khodarev, 2019), and accompanied IFN-inducible gene/protein expression (Figure 2C). The IPA Graphical Summary for the transcriptomic changes in astrocytes following F3-8 EV treatment is shown in Figure 3A (radial layout), with a very similar proteomic profile (Figure 3B). DDX58/RIG-I is the prominent central node connected to many pathways in stress and pathogen-sensing responses (and reduced activation of viral replication pathways). STRING analysis from the top 10 highly-expressed mRNAs from the transcriptome study shows tight associations of the OAS molecules with upregulated IFN-inducible genes (Figure 3C), and Biologic Pathways identified by the FunRich program indicate Type I and II IFN signaling (with a RIG-I/MDA5 basis), cytokine signaling in general, and immune responses (Figure 3D). Using a multiplex IFN ELISA for Type I, II, and III IFNs (and other inflammatory cytokines), we find that a variety of IFNs (as well as tumor necrosis factor alpha (TNFA), interleukin 1 alpha (IL1A), and chemokine (C-X-C motif) ligand 10 (CXCL10) are released by F3-8 EV-treated astrocytes in significant quantities compared to control (or no) treatments (Figure 3E). Curiously IL6 is strongly released following normal (astrocyte) EV or F3-8 EV treatments. Of note, prior secretome analysis showed that IFNG, multiple interleukins, CXCL10, and TNFA were all released at higher levels from F3-8 EV-treated astrocytes (Oushy *et al*., 2018).

**Figure 3:**
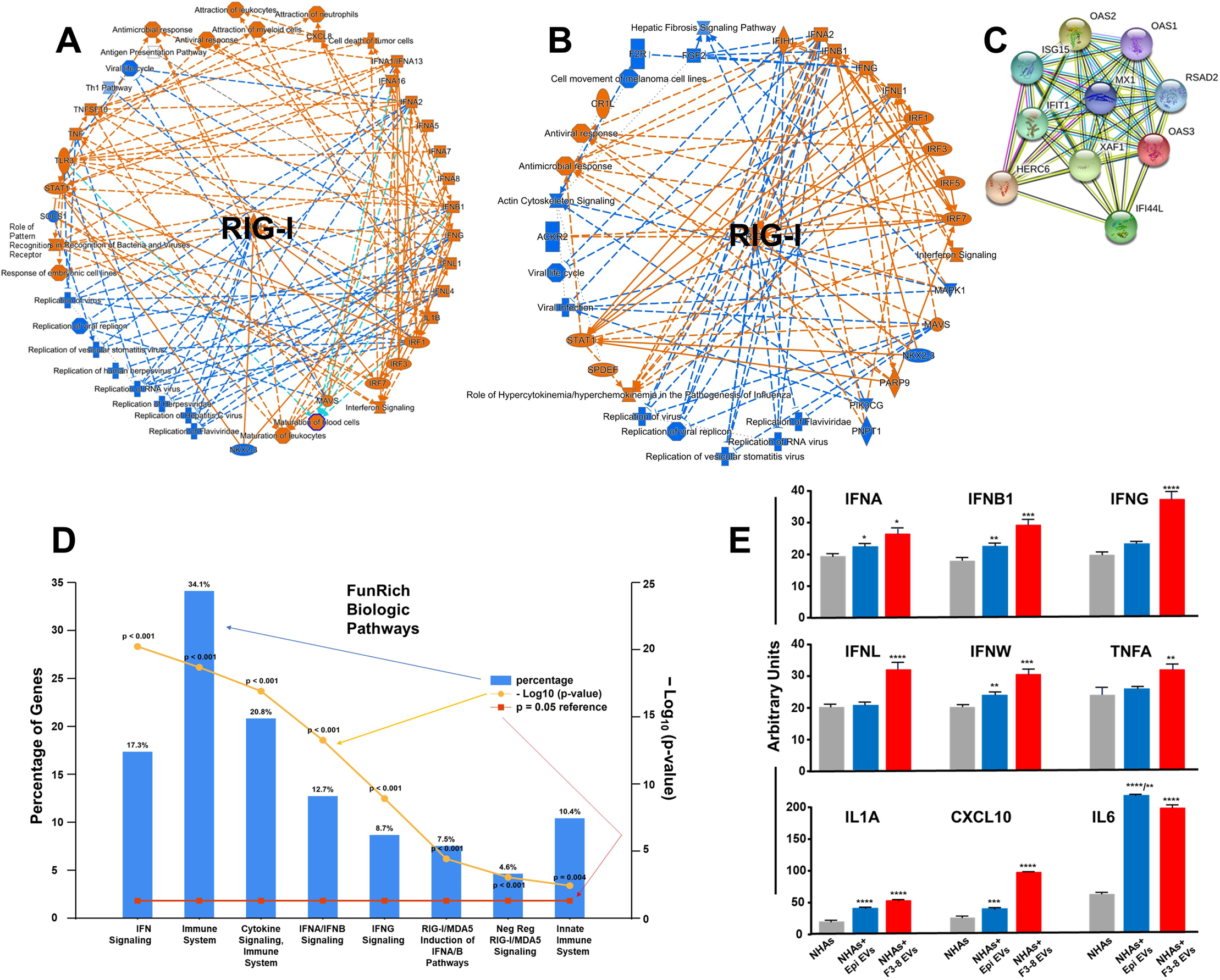
GBM F3-8 EVs induce an anti-viral response in recipient astrocytes. (A) Astrocyte transcriptomic analysis (IPA) following treatment with GBM F3-8 EVs and (B) proteomic analysis show RIG-I/DDX58 as the central nodes in the Graphical Summary (shown in radial layout). (C) STRING analysis of top 10 most highly over-expressed mRNAs in the astrocyte transcriptome following F3-8 EV treatment. (D) Top FunRich Biologic Pathways deduced from the astrocyte transcriptome following F3-8 EV treatment. (E) Following treatment with F3-8 EVs astrocyte supernatants were collected and subjected to ELISA analysis for type I-III interferons and other cyto/chemokines (PBL Assay Science VeriPlex Human Interferon 9-Plex ELISA kit cat # 51500). Gray bars = astrocytes (normal human astrocytes, NHAs) alone; blue bars = astrocytes treated with EVs from normal human epithelial cell (epi) EVs; red bars = astrocytes treated with F3-8 EVs. * p<0.05 compared to astrocytes alone; ** p<0.01 vs astrocytes alone; *** p<0.005 vs astrocytes alone; **** p<0.0001 vs astrocytes alone. For epi EV-treated astrocytes, IL6, /** p<0.01 vs F3-8 EV-treated astrocytes. One-way ANOVA followed by Tukey’s pairwise multiple comparisons.

We validated the upregulation of OAS1-3, MDA5/IFIH1, and RIG-I/DDX58 in Western blots where astrocytes were treated with a variety of tumor cell line EVs. Treated (or untreated) astrocytes were harvested, lysed, and lysates separated on SDS-PAGE followed by transfer to nitrocellulose and probing with antibodies shown (Figure 4). Consistent with transcriptomic and proteomic profiles, F3-8 EVs (and to some extent, F3-8 MVs/”microvesicles”) promoted expression of OAS1-3, MDA5/IFIH1, and RIG-I/DDX58 overall relative to EVs from other tumor cell lines, including G17-1 EVs. The IFN-inducible IFIT3 is also upregulated by F3-8 EVs/MVs. Interestingly, EVs from all the tumor cell lines induced higher GFAP expression in astrocytes compared to PBS control treatment, indicative of an activated – or perhaps reactive -- astrocyte state (Brenner, 1994; Sofroniew, 2014). We utilized the M2-7 and M6-7 cell line EVs because those tumors were recurrent and had received prior radiation therapy. There is a connection between tumor radiotherapy and stimulation of nucleic acid-sensing pathways (McLaughlin *et al*., 2020), and we wondered if that phenomenon might be transmitted to astrocytes by EVs from such tumors. This does not appear to be the case from this limited study.

**Figure 4:**
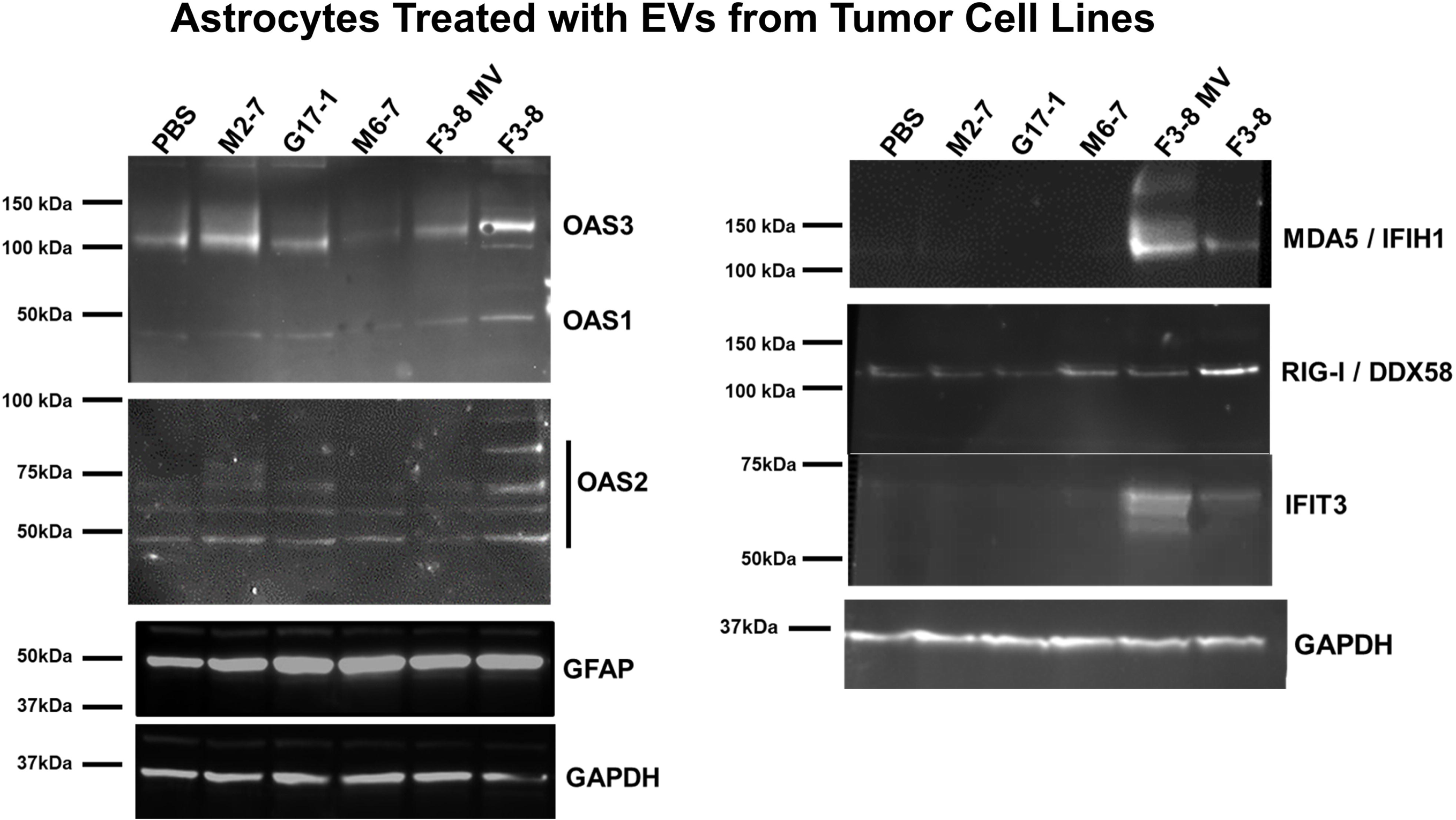
Western blot validation of innate immune/RNA sensors, IFN-induced molecules, and GFAP. EVs from cell lines shown (or PBS as control) were incubated with astrocytes for 24hr; cells were lysed, separated on SDS-PAGE, transferred to nitrocellulose, blocked and probed with antibodies against proteins listed, followed by washing and probes with secondary antibodies. That was followed by washing and chemiluminescent development. Molecular weight markers are indicated to the left of the blots. GAPDH was probed to verify comparable loading. Blots are shown as they appear in the FluorChem Q imager. “F3-8 MV”, astrocytes were treated with the same quantities of “microvesicles” derived from the F3-8 line (see Supplemental Figure 1 for details). M2-7 = adult (metastatic) embryonal rhabdomyosarcoma (recurrent; had prior radiation). M6-7 = Grade 4 astrocytoma, *IDH* mutant (recurrent; had prior radiation).

### 3.5. Astrocyte secretomes and signaling changes following treatment with G17-1 EVs

Turning to G17-1 EV-treated astroctyes and human brain slices, we examined the secretomes of brain tissues and astrocytes using a Proteome Profiler Cytokine Array, which displayed many changes in secreted molecules when comparing tissue or astrocytes treated with control EVs vs G17-1 EVs (Figure 5A, B). Entering these changes into IPA and employing the Comparison Analysis (Figure 5C) we note that there are evident similarities but also differences in the z-score-based hierarchical clustering of the top 25 Canonical Pathways. Certain elements of stress and cytokine signaling overlap, while other inflammatory pathways (e.g., IL8 signaling, chemokine signaling) appear relatively upregulated only in the brain secretome, and this is reflected in the heat maps (Figure 5A, B). The complexity of brain tissue may account for the differences compared to astrocytes alone, and the G17-1 EV-treated astrocyte secretome clearly differs from the F3-8 EV-treated astrocyte secretome (Oushy *et al*., 2018).

**Figure 5:**
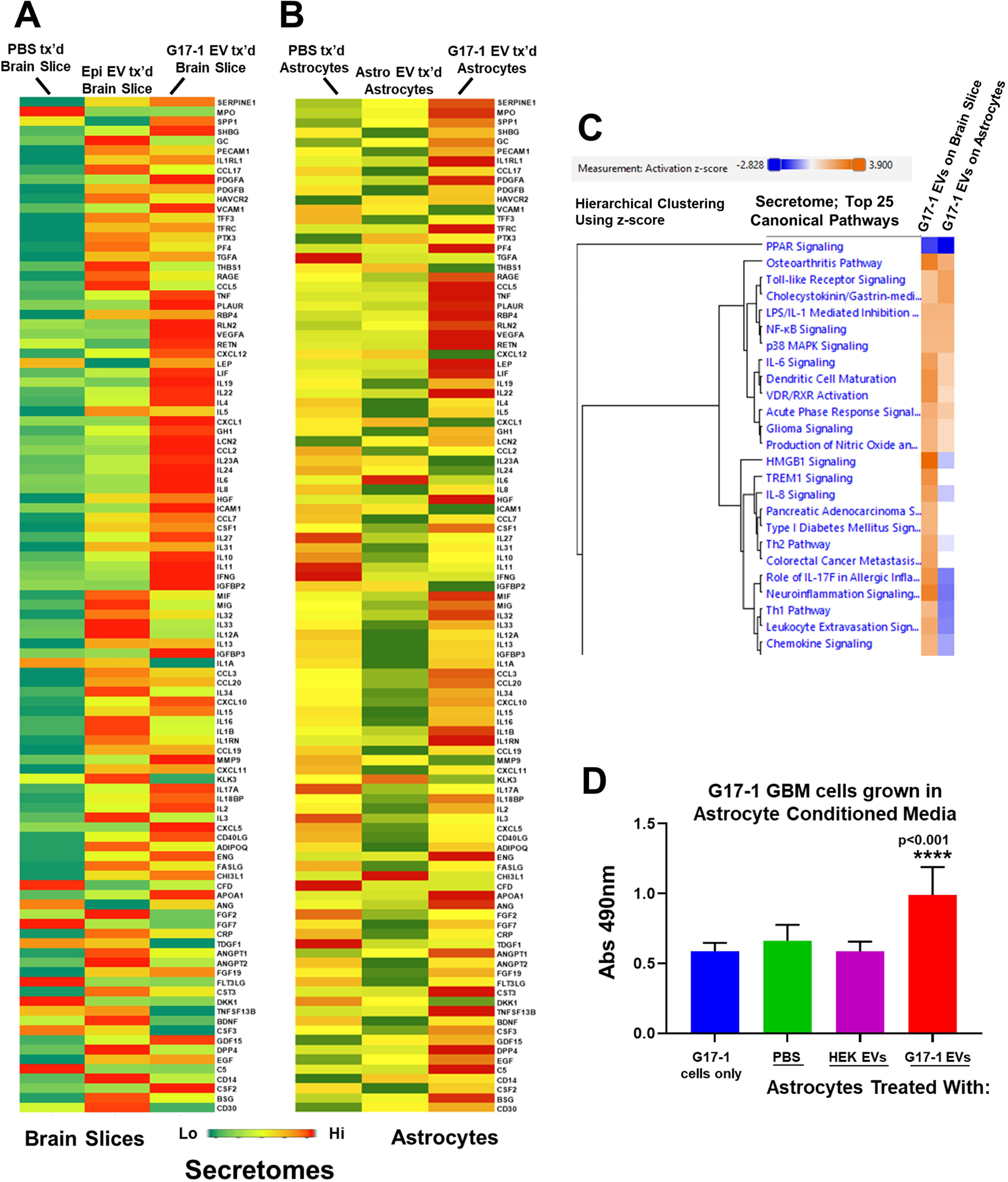
Brain slice and astrocyte secretomes following cell tissue and cell treatment with GBM G17-1 EVs. (A) Normal human brain slices were treated with PBS, normal epithelial cell EVs, or GBM G17-1 EVs. Conditioned media supernatants were subjected to Proteome Profiler Human XL Cytokine Arrays (ARY022B; R&D Systems). Spots were quantified by densitometry, averaged, and normalized to tissue weight. Results are displayed as heatmaps. (B) Astrocytes were treated with PBS, astrocyte EVs, or GBM G17-1 EVs. Conditioned media supernatants were subjected to Proteome Profiler Human XL Cytokine Arrays (ARY022B; R&D Systems). Spots were quantified by densitometry, duplicates averaged, and normalized to cell count. Results are displayed as heatmaps. (C) Using IPA Comparison Analysis, changes in cyto/chemokine expression of brain slices vs astrocytes (treated with G17-1 EVs) were categorized by Canonical Pathways as analyzed by hierarchical clustering by z-score. Top 25 Canonical Pathways compared by heatmaps are shown. (D) Astrocytes were treated with PBS, with EVs from HEK293 cells, or with G17-1 EVs. The conditioned media were transferred to G17-1 cells grown in the same ABM medium, and cell proliferation was measured by MTS assay 24hrs later.

We see increased release of numerous growth factors into astrocyte conditioned medium following G17-1 EV treatment. Based on this (and based on previous results using F3-8 EV-treated astrocyte conditioned medium (Oushy *et al*., 2018), we collected conditioned astrocyte medium following treatment of the astrocytes with G17-1 EVs and used it as a growth medium for G17-1 cells (Figure 5D). The conditioned medium promotes significantly increased proliferation rates for G17-1 cells compared to the typical growth of G17-1 or using astrocyte conditioned medium following HEK EV treatment. Despite the differences in the differentially-treated astrocyte secretomes (see (Oushy *et al*., 2018), both conditioned media promote growth of the respective cell lines.

IPA Network analyses on both the brain slice and astrocyte secretomes (following G17-1 EV treatment) identified networks where ERK1/2 were central nodes in radial layouts (Figure 6A, B), and ERK signaling was suggested in Figure 2E. We used a phospho-array to analyze changes in phosphorylation status of 43 targets on 39 different proteins (phospho sites are listed in Supplemental Table 4) in astrocytes treated with G17-1 EVs, and we modulated that by pre-treating the astrocytes with an ERK1/2 inhibitor before EV incubation. Notably, G17-1 EVs promote substantial changes (usually increases) in astrocyte protein phosphorylation status while ERK inhibition generally reduces that general phosphorylation phenotype (Figure 6C). However, there are some curious changes that show increased phospho status following ERK inhibition such as EGFR, mitogen-and stress-activated kinases 1 and 2 (MSK1/2), Ak strain transforming/protein kinase B (AKT) (T308), p53 (S392 and S46), focal adhesion kinase (FAK), and WNK (“with no lysine”) lysine deficient protein kinase 1 (WNK1). In astrocytes treated with ERK inhibitor, but not EVs, protein kinase AMP-activated catalytic subunit alpha 1/ PRKAA1 (AMPKA1), p53 (S392), STAT2 and 6, and platelet-derived growth factor receptor (PDGFR) all show increased phospho status, suggesting that ERK inhibition in either resting astrocytes or those treated with G17-1 EVs may lead to complicated signaling scenarios.

**Figure 6:**
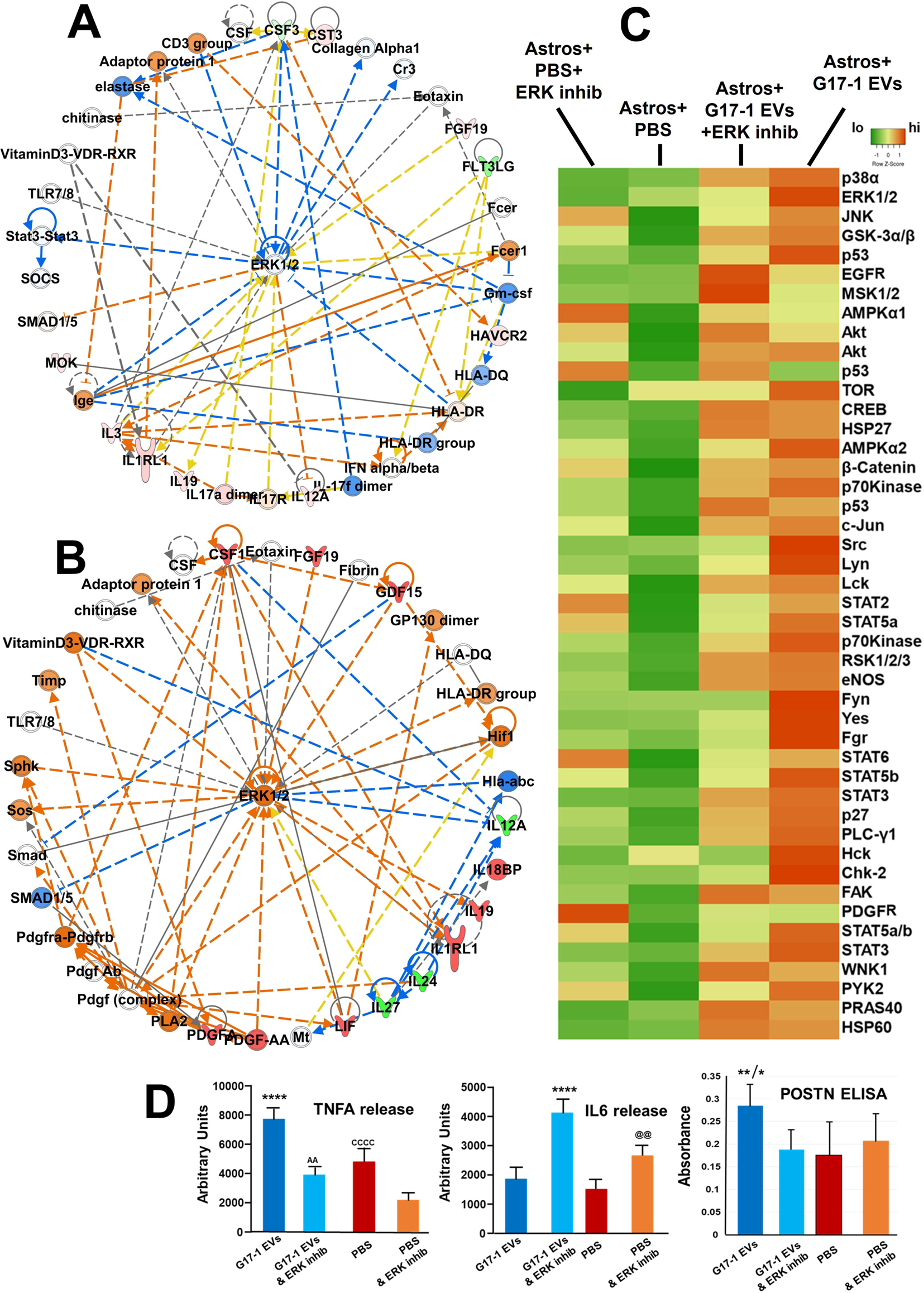
Brain slice and astrocyte secretomes following tissue and cell treatment with GBM G17-1 EVs implicate ERK1/2 signaling. (A) IPA network analysis of the secretome of brain slices treated with GBM G17-1 EVs identified a network with ERK1/2 signaling as the major node when presented in radial layout: “Cell Death and Survival; Cell Development; Inflammatory Response”. Score = 18, 10 Focus Molecules. (B) IPA network analysis of the secretome of astrocytes treated with G17-1 EVs identified a network with ERK1/2 signaling as the major node when presented in radial layout: “Cardiovascular System Development and Function; Hematological System Development and Function; Inflammatory Response”. Score = 18, 11 Focus Molecules. Score and Focus Molecule definitions are the same as in Figure 2. (C) Equal numbers of astrocytes were left untreated, or were treated with the ERK1/2 inhibitor SCH772984 (1μM) for 4hrs, and then +/- G17-1 EVs for 24hrs. Astrocytes were lysed, and lysates subjected to a Creative Biolabs Human Phospho-Kinase Antibody Array (AbAr-0225-YC). Spots were quantified by densitometry and duplicates averaged, and values were represented by heatmaps. (D) Astrocyte culture supernatants (from cells treated as in (C)) were subjected to the same ELISA as in Figure 3E; only TNFA and IL6 results are shown, along with a POSTN ELISA (ELH-POSTN; RayBiotech). For TNFA, **** p<0.0001 G17-1 EVs vs G17-1 EVs & ERKi; vs PBS; vs PBS & ERKi. AA p=0.0078 G17-1 EVs & ERKi vs PBS & ERKi. CCCC p<0.001 PBS vs PBS & ERKi. For IL6, **** p<0.0001 G17-1 EVs & ERKi vs G17-1 EVs; vs PBS; vs PBS & ERKi. @@ p=0.0012 PBS & ERKi vs G17-1 EVs. For POSTN, ** p<0.003 G17-1 EVs vs PBS; vs PBS & ERKi; * p<0.05 G17-1 EVs vs G17-1 EVs & ERKi. ANOVA followed by Tukey’s pairwise multiple comparisons.

The transcriptomes, proteomes, and secretomes of G17-1 EV-treated astrocytes all indicated upregulation and release of TNFA, IL6, and periostin (POSTN) compared to control settings, which we validated with either the same multiplex ELISA as used in Figure 3E or a specific POSTN ELISA (Figure 6D). ERK1/2 inhibition had curious effects in that TNFA and POSTN releases were indeed reduced upon ERK inhibition from G17-1 EV-treated astrocytes, but IL6 levels actually increased (even from astrocytes without EV treatment). This again suggests that ERK inhibition of astrocytes in the presence or absence of G17-1 EVs may have complicated outcomes.

### 3.6. Differential proteolytic effects of GBM

EVs themselves and after incubation with astrocytes Conditioned media from astrocytes treated with GBM EVs presented indications of extracellular protease release (dipeptidyl peptidase 4/DPP4 and the soluble plasminogen activator, urokinase receptor/PLAUR) in G17-1 EV-treated astrocyte conditioned media, Figure 5B; DPP4, MMP9, and soluble PLAUR in F3-8 EV-treated astrocyte conditioned media (Oushy *et al*., 2018). Using an extended protease array, we found that PLAUR and MMP2 are released by treated and untreated astrocytes, Cathepsin D is released by untreated and G17-1 EV-treated astrocytes, and MMP7 is released by astrocytes following G17-1 EV treatment. Cathepsin S, DPP4, and MMP9 shows slight elevations from G17-1 EV-treated astrocytes (Figure 7A). The protease array does not indicate protease activity, so we measured relative MMP (collagenase) activity directly in conditioned astrocyte media following treatment with F3-8, G17-1, M6-7, or M16-8 EVs. Curiously, only the G17-1 EV-treated astrocyte conditioned media showed significant collagenase activity (Figure 7B). As it is possible that all of the protease activity is in the G17-1 EVs used in the assay, we directly compared the G17-1 EV-treated astrocyte media with G17-1 EV protease activity. While there was significant activity from the EVs themselves, the conditioned media had significantly higher activity over most of the 1hr time course (Figure 7C). Curiously, F3-8 EVs themselves did indeed possess significant collagenase activity, while the media from astrocytes treated with those EVs showed minimal (background) activity when compared directly (Figure 7D). Utilizing brain slices as G17-1 EV recipients, we performed similar collagenase assays and again found activity in the conditioned media was greater than that of the EVs themselves, which was greater than controls (including slices treated with normal epithelial cell EVs at background levels; Figure 7E). Finally, using 2-photon microscopy, we see marked digested extracellular spaces in G17-1 EV-treated human brain slices (Figure 7F right panels) exhibiting morphological structural changes, while control slices (PBS treatment; epithelial cell EV treatment) show intact surfaces (Figure 7F left and middle panels). Top panels are single 2-D planes, while bottom panels are 3-D Z-stack reconstructions (“forest plots”). Oxygraph measurements confirm the tissue slices are viable (data not shown).

**Figure 7:**
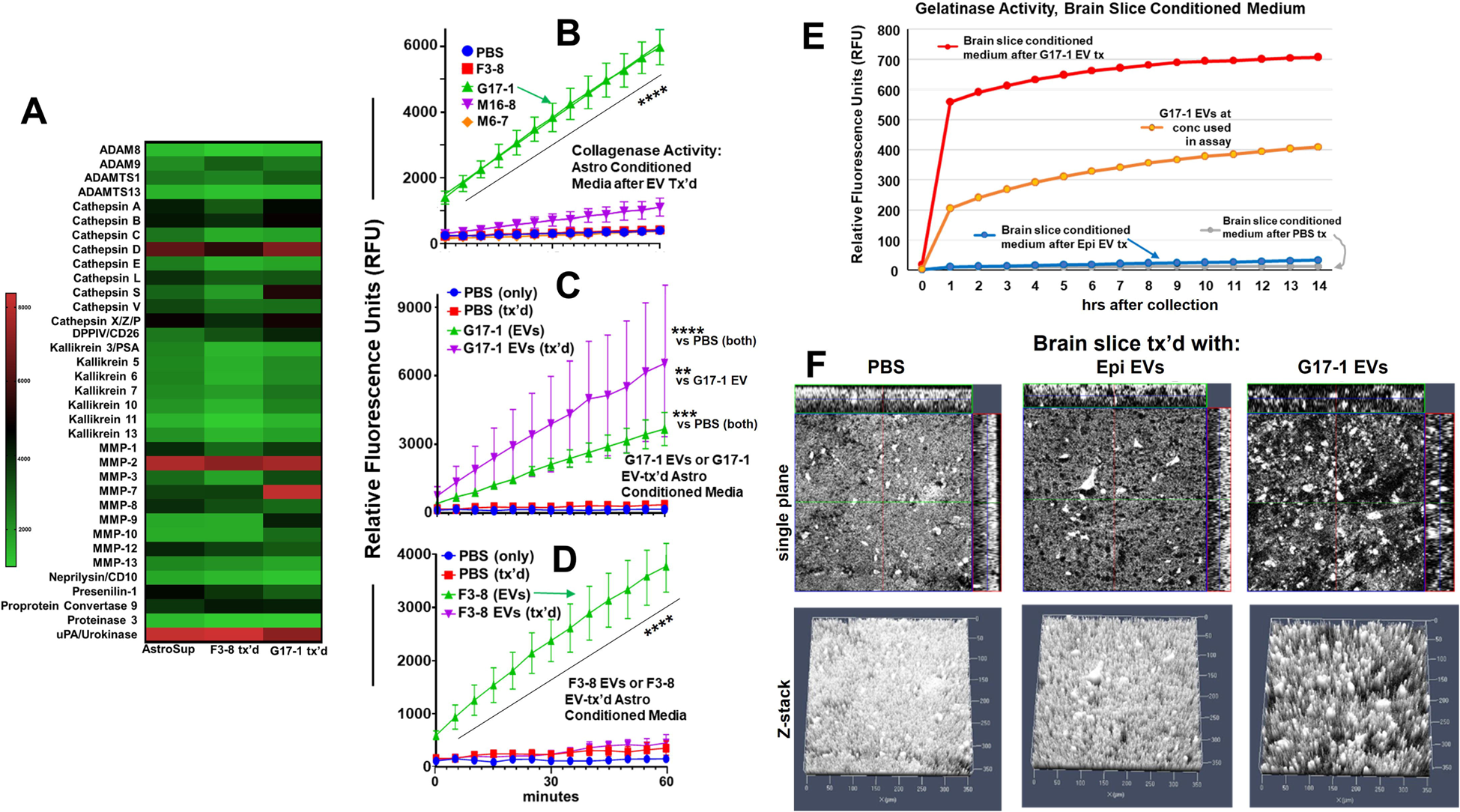
Proteases and activities in GBM EVs and astrocyte or brain slice conditioned media following GBM EV treatment. (A) Equal numbers of astrocytes were treated with PBS, GBM F3-8 EVs, or GBM G17-1 EVs for 24hrs. Conditioned media supernatants were collected and used to probe Proteome Profiler Human Protease Arrays (#ARY021B; R&D Systems). Spots were quantified by densitometry and duplicates averaged, and values were represented by heatmaps. (B) Collagenase/MMP activity was measured by abcam MMP Activity Assay Kits (Cat # ab112146); astrocytes were treated with the tumor EVs listed (PBS as a control; F3-8, G17-1, M16-8, M6-7) for 24hrs. Conditioned media were harvested, and assay over a 1hr period; **** p<0.0001 G17-1 EV treatment vs all others. (C) Collagenase/MMP activity of PBS only (blue line), astrocyte conditioned medium following PBS treatment (red line), G17-1 EVs only (green line, same concentration as used in astrocyte treatment), or astrocyte conditioned medium following G17-1 EV treatment (purple line). **** p<0.0001 G17-1 EV treatment of astrocytes vs either PBS; *** p<0.001 G17-1 EVs alone vs either PBS; ** p<0.01 G17-1 EV treatment of astrocytes vs G17-1 EVs alone. (D) Collagenase/MMP activity of PBS only (blue line), astrocyte conditioned medium following PBS treatment (red line), F3-8 EVs only (green line, same concentration as used in astrocyte treatment), or astrocyte conditioned medium following F3-8 EV treatment (purple line). **** p<0.0001 F3-8 EVs only vs all others. ANOVA followed by Tukey’s pairwise multiple comparisons. (E) Brain slices were treated G17-1 EVs (red line), epithelial cell EVs (blue line), or PBS (gray line) for 24hrs. Brain slice conditioned media was assayed in a gelatinase assay (EnzChek™ Gelatinase/Collagenase Assay Kit; cat # E12055; ThermoFisher) over 14hrs. G17-1 EV gelatinase activity was measured directly (orange line, same concentration as used in astrocyte treatment) over the same period. (F) Brain slices were treated as in (E) and were imaged in 2-photon excitation microscopy through 10mm depths to reveal matrix degradation (G17-1 EV treatment panels, right side). Top row = representative single plane, multiple deep focal plane imaging, 2-D maximum intensity projection image size = 1024 x 1024 pixel resolution; bottom row = axial scanning reconstructed Z-stack series, 3-D Intensity projection (XY image size x: 1024, y: 1024, Z: 33, 8-bit) was reconstructed from Z-stack (sample (x: 353.90 um, y:353.90 um, z: 10.65 um, 33 slides).

## 4. Discussion

The GBM tumor microenvironment is a source of therapeutic resistance (Goenka *et al*., 2021; Zhang *et al*., 2019b) and the GBM TME remains a focus of intense study (Bikfalvi *et al*., 2023; Faisal *et al*., 2022), including the roles of GBM EVs in various TME niches (Graner, 2019; Russo *et al*., 2022). As astrocytes are among the most prominent of cell types in the GBM TME, we focused on astrocyte molecular changes in response to incubation with GBM EVs. Our report here suggests that the roles of GBM EVs in affecting changes in astrocyte responses will vary depending on the tumor source of those EVs.

The heterogeneity of GBMs has long been considered a hallmark of the disease (Aum *et al*., 2014), even inferred in the older terminology “glioblastoma multiforme”. That different GBM EVs could have differential effects on cells of the TME should almost be expected. Nonetheless, in our hands, at least two outcomes – an activated/reactive astrocyte status, and tumor cell growth in response to EV-treated astrocyte conditioned media – remain consistent features no matter which GBM EVs are used (Figures 4, and 5, and (Oushy *et al*., 2018). This implies there may be multiple means to the same ends. The transcriptomic and proteomic analyses (Figures 1, 2; Supplemental Figures 2-6) of GBM EV-treated astrocytes painted broad strokes of differential molecular phenotypes induced by the two different spheroid line EVs that extended to the astrocyte secretomes and activities therein (Figures 3, 5-7).

A consistent theme within astrocyte responses to F3-8 GBM EVs included the interferon-related anti-viral responses. Intertwined within the type I and type II IFN pathways (either driven by IFNs or leading to IFNs, Supplemental Figure 8A) are interleukin pathways that play into each other (Supplemental Figure 8A, B). The noted role of RIG-I/DDX58 (along with MDA5/IFIH1) as a central node in these inflammatory pathways (Figure 3A,B) could be driven by RNAs from EVs, but was only seen with F3-8 GBM EV treatment. The mechanisms behind this await further study. The nucleic acid sensors downstream connect their innate immune roles to nuclear factor kappa B (NFKB) complex activation (De Miranda *et al*., 2009; Furr *et al*., 2010), which further ties to adaptive immune responses.

From the transcriptomics of astrocytes treated with F3-8 EVs, we see the immune response networks related to NFKB complex activation in Supplemental Figure 9A, leading to a putative accentuated major histocompatibility complex (MHC) I antigen presentation pathway (such pathways are among the top Canonical Pathways in Supplemental Figure 3). Paradoxically, there appear to be reduced quantities of MHC II component expression, suggesting reduced antigen presentation to CD4+ T cells (Supplemental Figure 9B). Astrocytes are historically known to be potential antigen presenting cells (Fierz *et al*., 1985), and the transcripts for immune synapse members such as some MHC I molecules and co-stimulatory molecules CD40, inducible T cell co-stimulator ligand (ICOSLG), and ICAM1 are significantly elevated in F3-8 EV-treated cells (Supplemental Table 1). However, immune checkpoint molecules CD274/programmed cell death ligand 1 (PD-L1) and programmed cell death 1 ligand 2 (PDCD1LG2)/(PD-L2) and other inhibitors (indoleamine 2, 3-dioxygenase 1/IDO1) are also significantly upregulated, which is a known effect of IFN in the TME (Fierz *et al*., 1985; Gruosso *et al*., 2019). Astrocyte effectiveness at initiating T cell responses may be contextual, or even protective in limiting immune responses (Gimsa *et al*., 2013); curiously, murine astrocytes in a viral infection scenario upregulate both MHC II and PD-L1 in a suppressive response to IFNG (Schachtele *et al*., 2014). The shift away from MHC II antigen presentation along with the expression of immune inhibitors suggests the generation of an immune suppressed environment following F3-8 EV incubation, which benefits the tumor. This may be despite the attempts to increase antigen presentation via MHC I pathways, implying that the astrocytes attempt to inhibit tumor progression, but ultimately are co-opted. Further studies are needed to assess this situation.

Astrocytes treated with G17-1 GBM EVs showed consistent signs of cell-cell communication via junctional signaling in proteomic and transcriptomic analyses (Figure 2D; Supplemental Figures 4 and 6). The extracellular space was also an area of importance. The genes encoded by the top 10 highly-expressed mRNAs are all either membrane/extracellular matrix proteins or are secreted outside the cell, most with network connections to each other and with gene ontology assignments to the extracellular space (STRING analysis, Supplemental Figure 10A, B). Extracellular proteolytic activity was evident not only in G17-1 EVs (and also F3-8 EVs, Figure 7B), but also in G17-1 EV-treated astrocyte and brain slice conditioned media (Figure 7B, E, F). Such proteolysis at two levels implies that the G17-1 GBM could utilize both its own EVs and the astrocytes those EVs encounter to alter the extracellular space, allowing for more efficient tumor cell migration/invasion. Given that the G17-1 EV-treated astrocyte conditioned media was perhaps more proteolytically active than the GBM EVs themselves, this suggests an amplification of the input. The nature of this response is under current study. Curiously, EVs from other tumors did not induce astrocyte extracellular MMP activity (Figure 7B), again emphasizing the singular character of a given GBM and its EVs. These data contrast with astrocyte gelatinase activity seen with GBM EV-treated astrocytes in direct contact with gelatin substrates (Hallal *et al*., 2019), where different GBM EVs promoted differing levels of astrocyte gelatinase. In our experiments, we examined only the secreted protease activity which differentially associated only with G17-1 EV treatments (of the four examined); further studies are needed to clarify the extent of this response.

Reactive astrocytes for decades have been described as components of brain tumors (Tascos *et al*., 1982; Vinores and Rubinstein, 1985), and their contributions to GBM malignancy are expanding with further study (Guan *et al*., 2018). There are numerous ways by which reactive astrocytes may aid in GBM progression, including various secreted factors (Matias *et al*., 2018) but also involving direct contact via gap junctional communications (Zhang *et al*., 1999). Importantly, the metabolic effect of GBM EVs on astrocytes is getting recognition (Zeng *et al*., 2020) as part of the larger metabolic impact reactive astrocytes have in the TME (Zhang *et al*., 2020).

The roles of GBM EVs in generating reactive astrocytes are poorly understood (Nieland *et al*., 2021), but EVs may be seen as literal extensions of the tumor. The elevated expression of GFAP in EV-treated astrocytes compared to untreated controls (Figure 4) may be taken as a sign of a conversion to “reactive astrocyte” status. Given the difficulty in determining differences between “active” astrocytes vs “reactive” astrocytes vs “astrogliosis”, we employ the literature’s consensus term “reactive astrocyte” for such changes due to pathologic conditions (Escartin *et al*., 2021). That publication notes the widely varying concepts of astrocyte reactivity, the different pathologies that may drive heterogenous populations of astrocytes to respond in different ways, and the putative markers described that may need to be incorporated into larger signatures of “reactive astrocytes”. Our data fit piecemeal into these markers and categories; for instance, glial cell line-derived neurotrophic factor (GDNF) mRNA is significantly upregulated in G17-1 EV-treated astrocytes, but not F3-8 EV-treated astrocytes (Supplemental Table 1). However, the NFKB signaling (Figure 5, Supplemental Figure 9A), secretome changes (Figure 5, and (Oushy *et al*., 2018), STAT3 signaling (Figure 6C and (Oushy *et al*., 2018), and mitochondrial/metabolic changes (Paucek et al, manuscript in preparation) reflect reactive astrocyte phenotypes. We also note that transcriptomic and proteomic analyses find Canonical Pathways relating to hepatic stellate cell activation and hepatic fibrosis signaling pathways (Supplemental Figures 2 and 3). Hepatic stellate cells are fibrotic responders in the liver and produce GFAP when activated; they display other pathways congruent with astrogliosis, including release of hepatocyte growth factor (HGF) (Figure 5A, B; (Oushy *et al*., 2018; Schachtrup *et al*., 2011). This suggests overlap in general patterns of stimulated reactive cell types in protective circumstances.

We point out that all the astrocytic changes here result solely from treatment with GBM EVs; there is never physical interaction with the GBM cells themselves. This begs the question of what triggers do the EVs provide to stimulate the astrocytes? Extracellular promoters or regulators of reactive astrocytes/astrogliosis include high mobility group box 1 (HMGB1) and IL6 (Sofroniew, 2014), which are at higher content on F3-8 EVs vs G17-1 EVs (Graner et al, manuscript in preparation). Low quantity entities such as cytokines and chemokines may be within the EV lumen, and/or on EV surfaces, and likely require greater sensitivity assays than shotgun proteomics (Fitzgerald *et al*., 2018; Fringuello *et al*., 2021). HMGB1 as a secreted “danger signal”/alarmin has neuroinflammatory capabilities (Frank *et al*., 2015) and is known to form complexes with other inflammatory cytokines (Yang *et al*., 2020). HMGB1 can be trafficked via EVs (Murao *et al*., 2021; Yang *et al*., 2020), including rat glioma EVs (Ma *et al*., 2019), and one may speculate that vesicles laden with alarmins and possibly pro-inflammatory cytokines could induce inflammatory responses in recipient astrocytes, followed by further autocrine stimulation. The reactive astrocyte milieu presumably benefits the tumor.

There is one recurring overall question regarding GBM EV pro-tumor impacts on the TME: can we mitigate these impacts therapeutically? GBM EVs clearly affect astrocytes in ways that ultimately appear to aid the tumor in terms of survival and resistance, progression, and invasion (Bian *et al*., 2019; Colangelo and Azzam, 2020; Gao *et al*., 2020; Hallal *et al*., 2019; Ma *et al*., 2019; Oushy *et al*., 2018; Taheri *et al*., 2018; Yu *et al*., 2018; Zeng *et al*., 2020). Our work here confirms and extends this information to imply that EVs from different GBM cell lines promote different molecular phenotypes in recipient astrocytes. This suggests that targeting specific aspects or pathways based on a particular model may be inadequate to prevent EV-driven protumor events when expanded to tumors beyond that model. A truly generalized method to prevent EV-driven tumor enhancement would be to specifically prevent tumor EV release. It is not clear that such specific inhibitors exist despite substantial efforts to find them (Catalano and O’Driscoll, 2020; Rezaie *et al*., 2022), as repressing the various modes of EV biogenesis and specificity towards cancer cells remain difficult barriers. Another area for EV interruption could be in preventing EV/target cell interactions. We recently identified phage-display peptides reactive to GBM EV surfaces that prevented complement-dependent neurotoxicity generated by GBM EVs (Zhou *et al*., 2022a; Zhou *et al*., 2022b). However, these peptides are also GBM EV-specific, implying that specificity is possible, but that multiple epitopes would need to be targeted for generalized, “off-the-shelf” coverage (even without delivery considerations). These are among the many variables necessarily accountable in terms of targeting tumor EVs in the TME.

## 5. Conclusion

GBM EVs affect cells of the TME; in GBMs, many of those cells are as astrocytes, based on tumor origin and brain cellular content. The TME may initially resist GMB growth and progression, but ultimately is co-opted to support the tumor. GBM EVs dramatically impact TME cells, and that impact differs depending on the GBM origin of the EVs. We show here that EVs from two different GBMs drove transcriptomic, proteomic, and secretomic changes in astrocytes and brain tissues that varied in molecular outcomes and functional readouts, but still drive reactive astrocyte status and produce a tumor-growth supportive milieu. Thus, there may be multiple means to the same end. This implies a “personalized” response by cells of the TME to different GBM EVs, and suggests that we recognize the limits of our individual model systems.

## Acknowledgements

The authors thank the CU Dept of Neurosurgery Nervous System Biorepository (https://medschool.cuanschutz.edu/neurosurgery/research-and-innovation/services/nervous-system-biorepository) for access to samples and cell lines, and to the CU Anschutz/CUACC Exosome Core for use of the Nanosight NTA and general assistance. We thank Dr. Anza Darehshouri, of the Electron Microscopy Core Facility (Department of Cell and Developmental Biology, University of Colorado Anschutz School of Medicine) for help with electron microscopy, Dr. Monika Dzieciatkowska of the Mass Spectrometry Proteomics Shared Resource Facility (RRID: SCR_021988), and the Genomics Shared Resource (RRID: SCR_021984), acknowledging the Cancer Center Support Grant (P30CA046934). We thank the Advanced Light Microscopy Core (ALMC) in the CU Neurotechnology Center for use of the 2-photon DIVER microscope. Mary Wang is a member of the University of Colorado School of Medicine Mentored Scholarly Activity program. Bryne Knowles, Charlotte McRae, and Vince Bolus were supported by the University of Colorado Cancer Center’s Cancer Research Experience for Undergraduates (CREU) from NIHR25CA240122 and UCCC. Brooke Metzer and Morgan Lenz were sponsored by the CU Dept of Neurosurgery/Illinois College Alumni Fellowship. The study was funded by NIH grants NIMH1R21MH118174-01and 4R33MH118174, and the Department of Neurosurgery Research Funds.

---

**Supplemental Figure 1:** GBM cell lines used in the study, and extracellular vesicle production and characterization. (A) F3-8 (left) and G17-1 (right) spheroids grown under stem cell-like conditions. Scale bars = 200mm for F3-8 image, 100mm for G17-1 image. (B) Schematic of steps involved in EV isolation. (C) Nanosight nanoparticle tracking analysis (NTA) of F3-8 EVs (Stats: Merged Data; Mean:110.6nm, Mode: 101.3nm, SD: 25.5nm), of G17-1 EVs (Stats: Merged Data; Mean: 135.8nm, Mode: 131.3nm, SD: 34.9nm), and of HEK293 EVs (Stats: Merged Data; Mean: 144.4nm, Mode: 126.9nm, SD: 33.9nm) with concentrations listed. (D) Transmission electron microscopy of F3-8 EVs (left, top, and two beneath) and G17-1 EVs (right panels) with scale bars/magnification listed on each image. (E) ExoCheck arrays showing putative typical EV markers for lysed EVs from cell lines: F3-8; G17-1; M2-7; M6-7; M16-8. The template is on the far right. (F) ExoCET (acetylcholinesterase) assay showing enzymatic activity and putative quantification for HEK, G3-8, and G17-1 EVs.

**Supplemental Figure 2:** Transcriptomic data volcano graphs for astrocytes treated with GBM EVs. Astrocytes were treated with GBM F3-8 EVs (left panel) or G17-1 EVs (right panel). Control treatment (“tx”) was with human epithelial (Epi) cell EVs. Astrocytes were extracted and the transcriptomes were analyzed on Human Clariom D chips; volcano graphs were generated in Transcriptome Analysis Console (TAC) 4.0.1 (ThermoFisher). Red dots show RNAs significantly upregulated with GBM EV treatments, while green dots show RNAs significantly upregulated with epithelial EV treatments.

**Supplemental Figure 3:** Ingenuity Pathway Analysis (IPA) modified summary reduced to 1 page. IPA modified summary for transcriptomics of astrocytes treated with GBM F3-8 EVs.

**Supplemental Figure 4:** IPA modified summary reduced to 1 page. IPA modified summary for transcriptomics of astrocytes treated with GBM G17-1 EVs.

**Supplemental Figure 5:** IPA modified summary reduced to 1 page. IPA modified summary for proteomics of astrocytes treated with GBM F3-8 EVs.

**Supplemental Figure 6:** IPA modified summary reduced to 1 page. IPA modified summary for proteomics of astrocytes treated with GBM G17-1 EVs.

**Supplemental Figure 7:** Ingenuity Pathway Analysis (IPA) network legends, shapes, and edge descriptions.

**Supplemental Figure 8:** Relevant IPA-derived networks from transcriptomics, astrocytes treated with GBM F3-8 EVs. (A) Network 3: “Antimicrobial Response, Infectious Diseases, Inflammatory Response”; Score = 38; Focus Molecules = 25. (B) Network 1: “Cell Cycle, Cell-to-Cell Signaling and Interaction, Cellular Development”; Score = 41; Focus Molecules = 26. Score and Focus Molecule definitions are the same as in Figure 2.

**Supplemental Figure 9:** Additional relevant IPA-derived networks from transcriptomics, astrocytes treated with GBM F3-8 EVs. (A) Network 4: “Cell Morphology, Hematological System Development and Function, Infectious Diseases”; Score = 32; Focus Molecules = 22. (B) Network 5: “Endocrine System Disorders, Gastrointestinal Disease, Immunological Disease”; Score = 26; Focus Molecules = 19. Score and Focus Molecule definitions are the same as in Figure 2.

**Supplemental Figure 10:** STRING analysis of the top 10 upregulated mRNAs from transcriptome of astrocytes treated with GBM G17-1 EVs. (A) STRING functional interaction protein network of the top 10 most highly upregulated mRNAs (compared to epithelial cell EV-treated astrocytes) from astrocytes treated with G17-1 EVs. (B) Gene Ontology terms relevant to the mRNAs in (A).

**Supplemental Table 1:** Transcriptome Analysis Console (TAC) 4.0.1 of astrocytes treated with GBM F3-8 EVs.

**Supplemental Table 2:** Transcriptome Analysis Console (TAC) 4.0.1 of astrocytes treated with GBM G17-1 EVs.

**Supplemental Table 3:** Proteomic data, astrocytes treated with GBM F3-8 or G17-1 EVs, or PBS.

**Supplemental Table 4:** Phosphorylation sites on proteins in Creative Biolabs Human Phospho-Kinase Antibody Array AbAr-0225-YC.

## Notes

### Competing Interest Statement

The authors have declared no competing interest.

