## Supplemental Table 4 for "A tale of two tumors: differential, but detrimental, effects of glioblastoma extracellular vesicles (EVs) on normal human brain cells"

Akt 1/2/3 (S473)

**Supplemental Table 4**

**Phosphorylation Sites on Proteins**

**in Creative Biolabs Human Phospho-Kinase Antibody Array AbAr-0225-YC**

Hck (Y411)

PLC gamma-1 (Y783)

Akt 1/2/3 (T308)

HSP27 (S78/S82)

PRAS40 (T246)

AMPK alpha1 (T183)

HSP60, Pyk2 (Y402)

AMPK alpha2 (T172)

JNK 1/2/3 (T183/Y185, T221/Y223)

RSK1/2/3 (S380)

beta-Catenin, Lck (Y394)

Src (Y419)

Chk-2 (T68)

Lyn (Y397)

STAT2 (Y689)

c-Jun (S63)

MSK1/2 (S376/S360)

STAT3 (S727)

CREB (S133)

p27 (T198)

STAT3 (Y705)

EGFR (Y1086)

p38 alpha (T180/Y182)

STAT5a (Y699)

eNOS (S1177)

p53 (S15)

STAT5a/b (Y699)

ERK1/2 (T202/Y204, T185/Y187)

p53 (S392)

STAT5b (Y699)

FAK (Y397)

p53 (S46)

STAT6 (Y641)

Fgr (Y412)

P70 S6 Kinase (T389)

TOR (S2448)

Fyn (Y420),

p70 S6 Kinase (T421/S424)

WNK-1 (T60)

GSK-3 alpha/beta (S21/S9)

PDGF R beta (Y751)

Yes (Y426)
