## Supplementary figures and images for "A tale of two tumors: differential, but detrimental, effects of glioblastoma extracellular vesicles (EVs) on normal human brain cells"

### Supplementary Figure 1

#### Slide 1
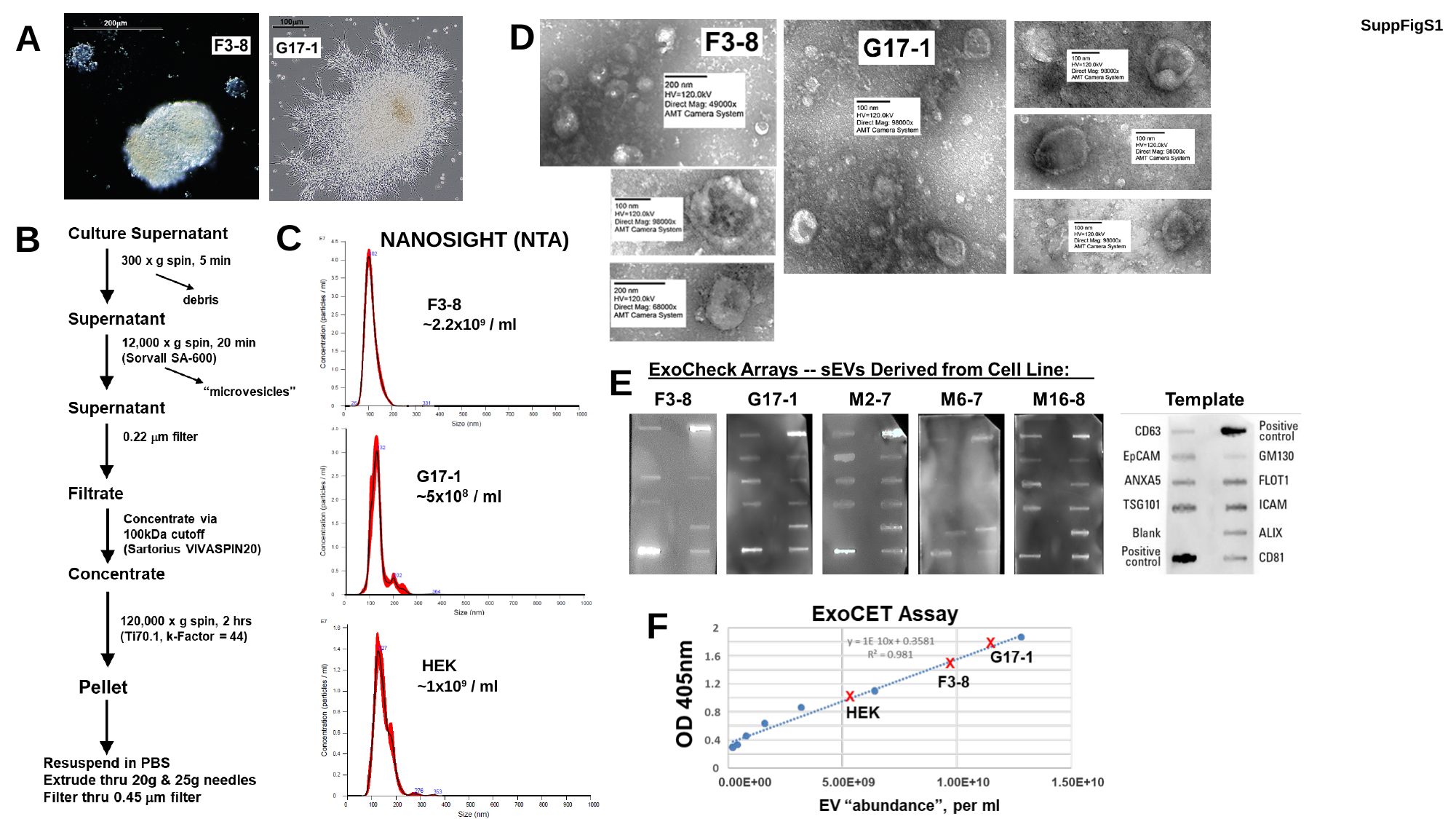

D
A
SuppFigS1
C
B
NANOSIGHT (NTA)
F3-8
~2.2x10^9^ / ml
E
F
HEK
~1x10^9^ / ml

### Supplementary Figure 2

#### Slide 1
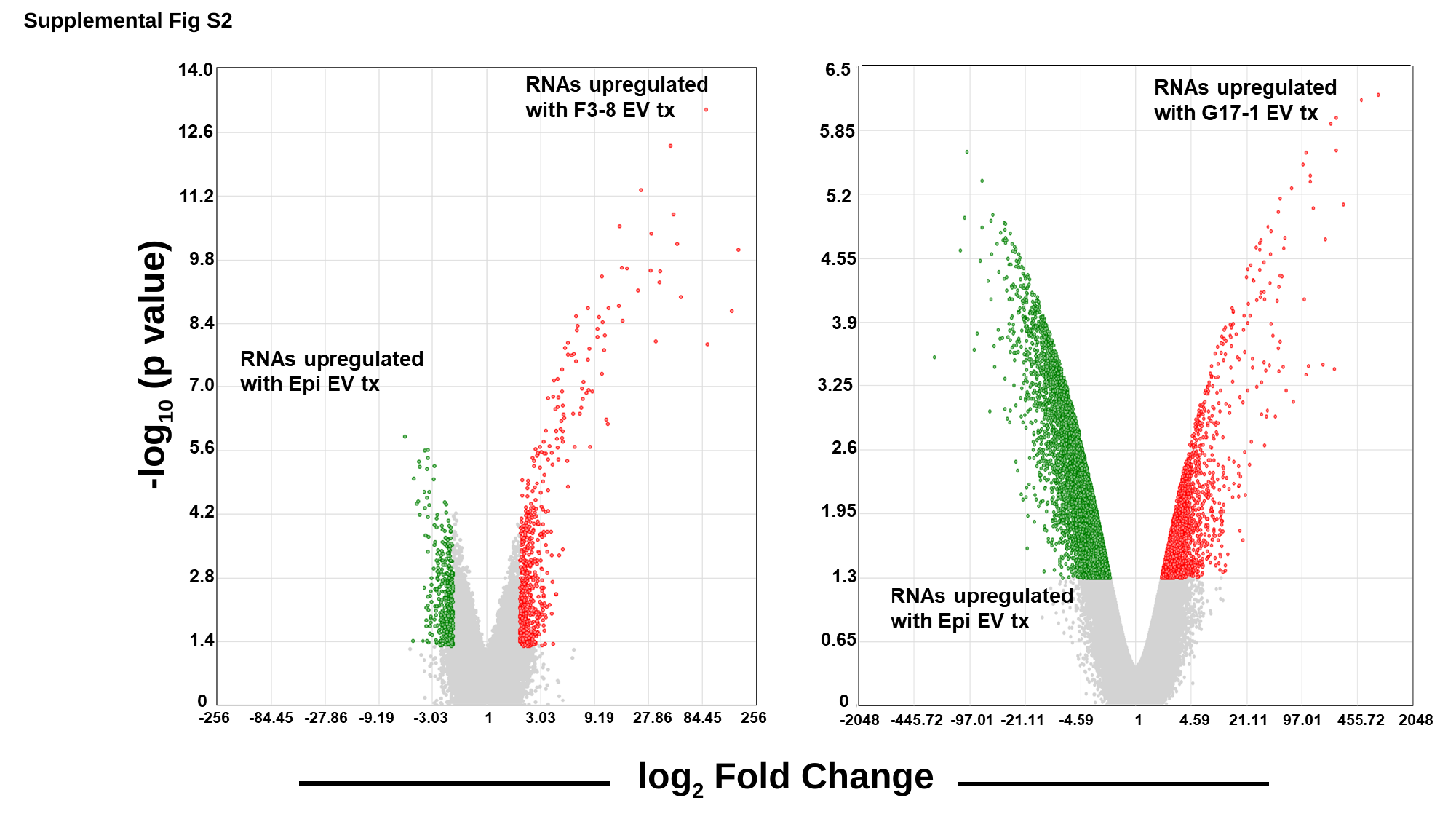

Supplemental Fig S2
-log_10_ (p value)
log_2_ Fold Change

### Supplementary Figure 10

#### Slide 1
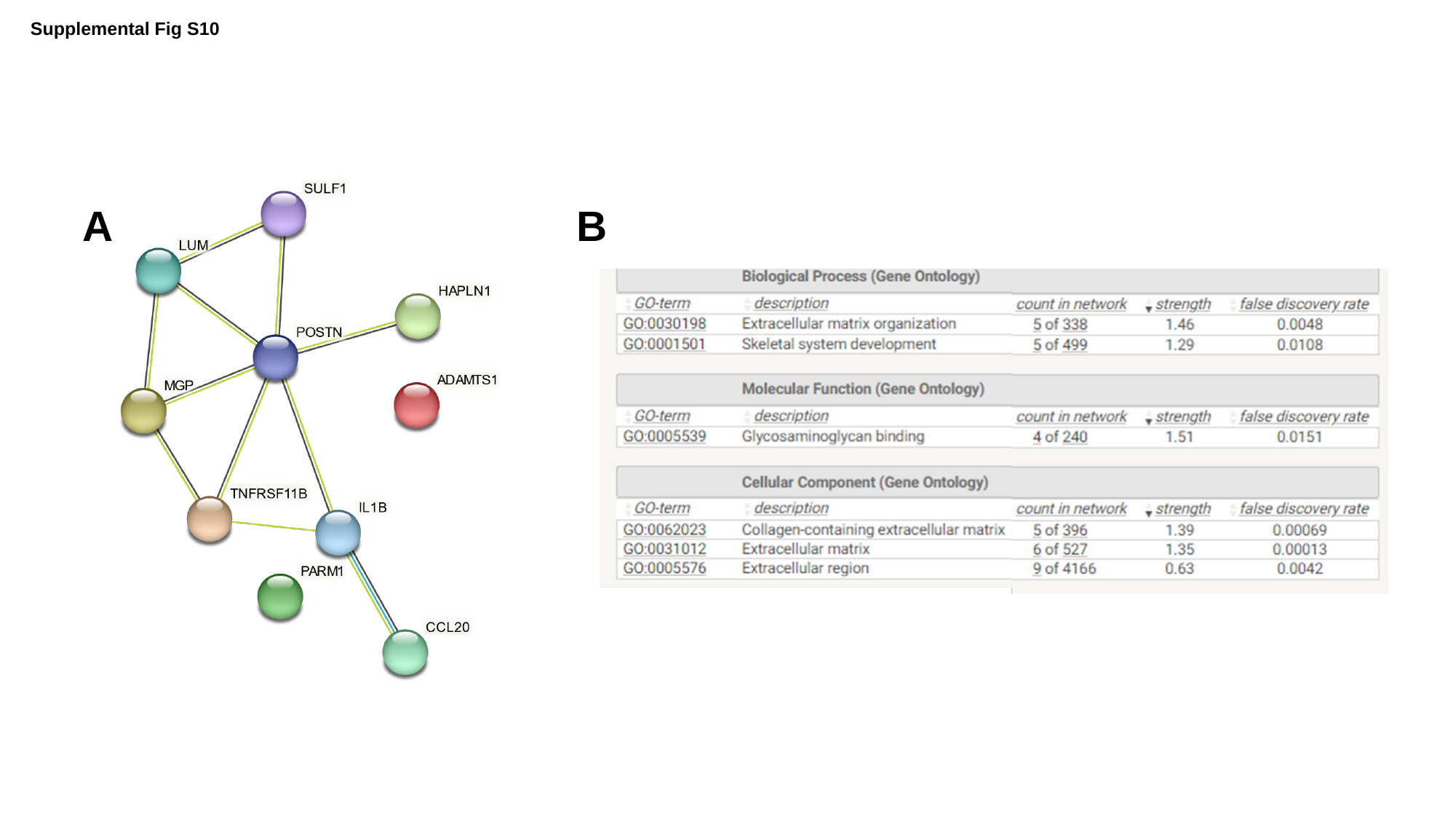

Supplemental Fig S10
A
B
