## Supplementary Figure 3,4,5,6 for "A tale of two tumors: differential, but detrimental, effects of glioblastoma extracellular vesicles (EVs) on normal human brain cells"

### Slide 1
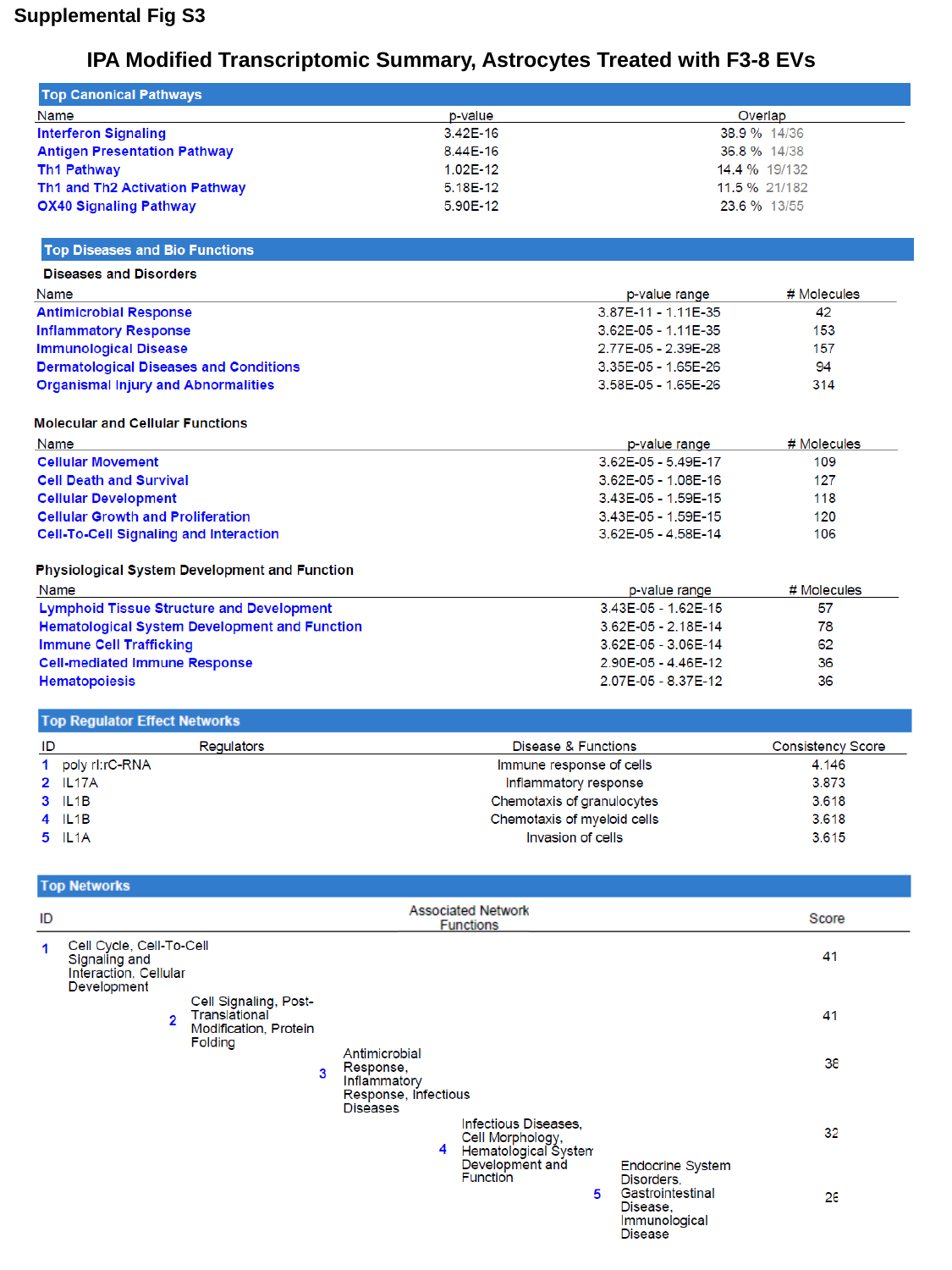

Supplemental Fig S3
IPA Modified Transcriptomic Summary, Astrocytes Treated with F3-8 EVs

### Slide 2
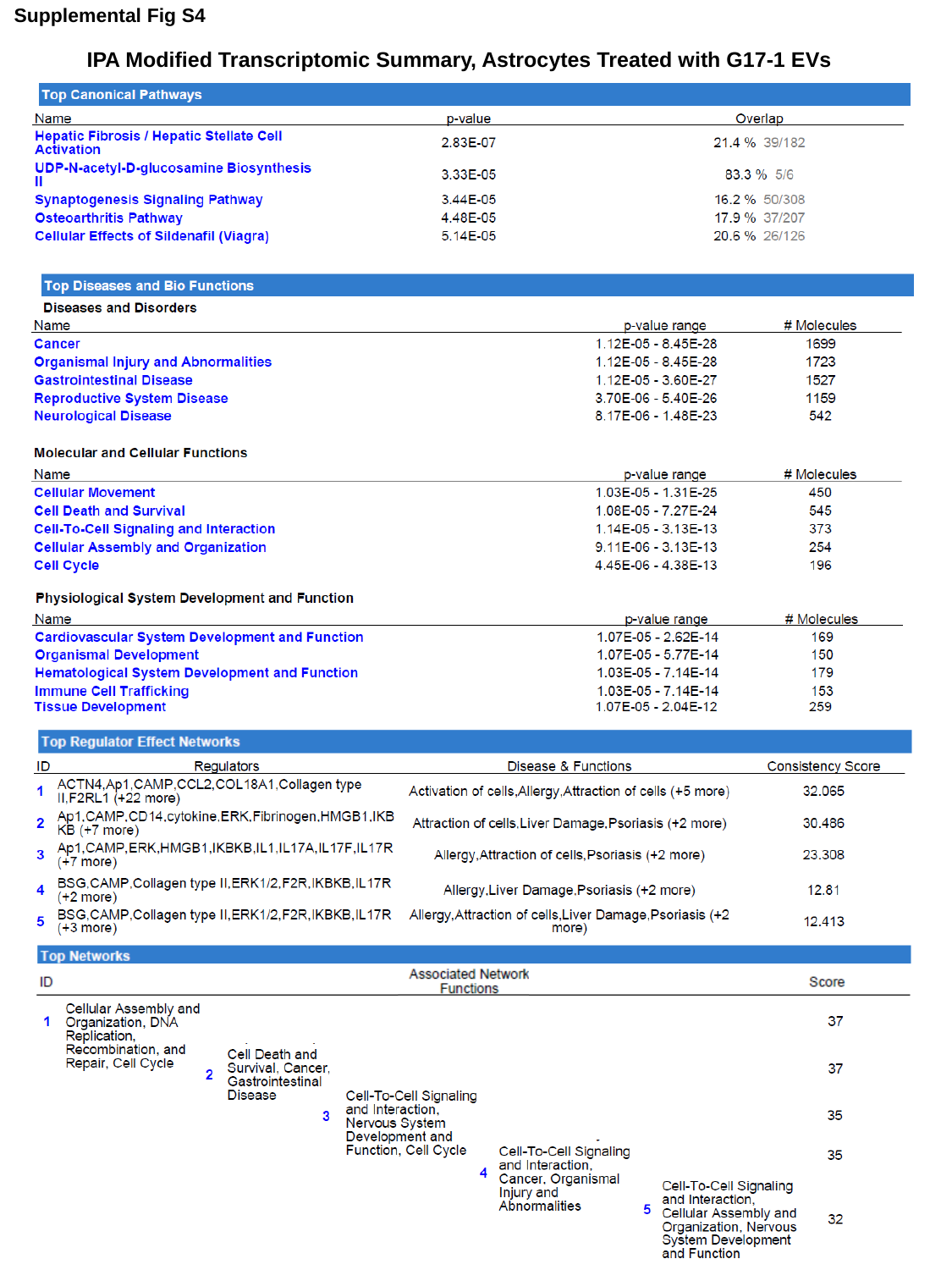

Supplemental Fig S4
IPA Modified Transcriptomic Summary, Astrocytes Treated with G17-1 EVs

### Slide 3
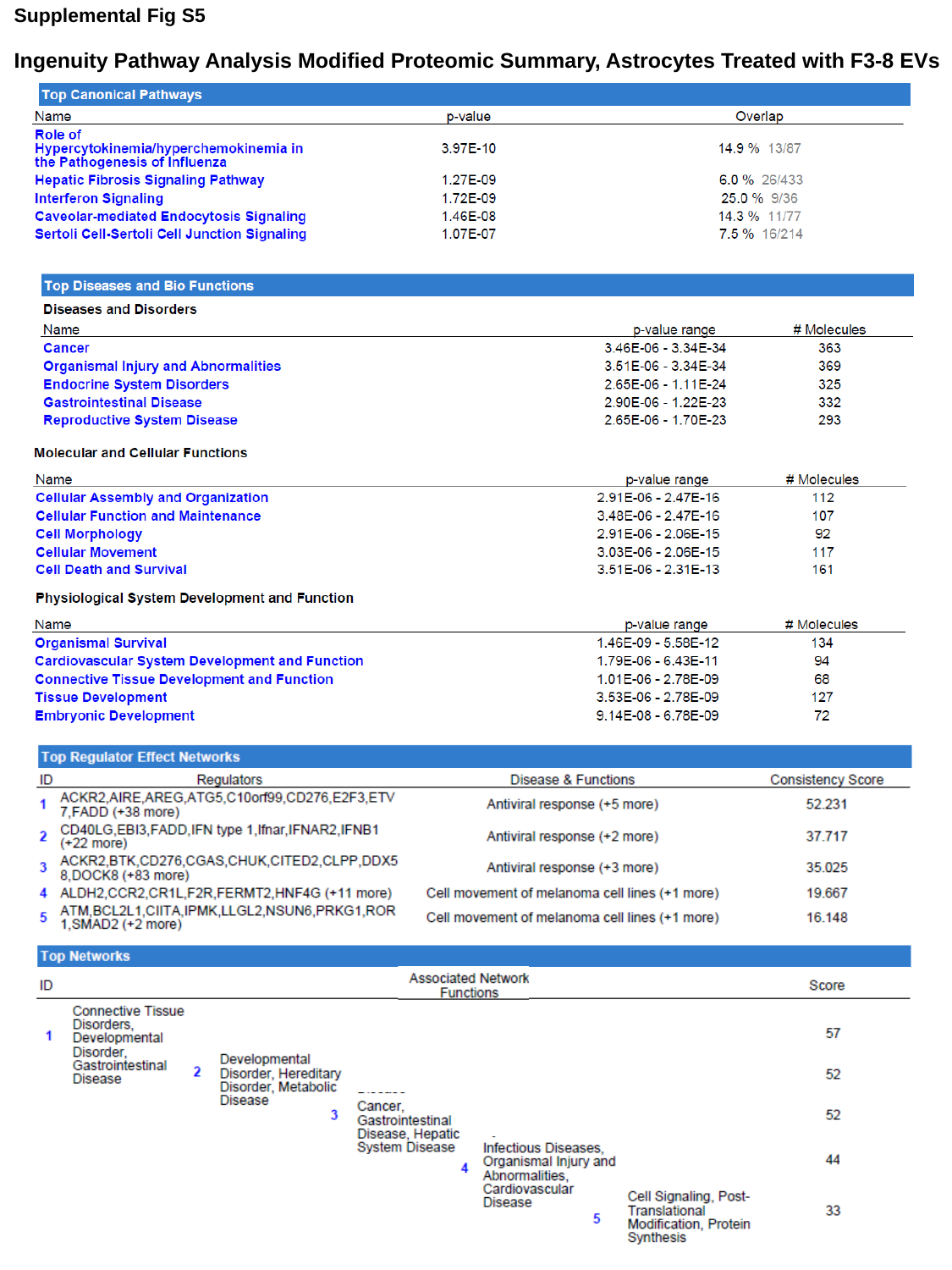

Supplemental Fig S5
Ingenuity Pathway Analysis Modified Proteomic Summary, Astrocytes Treated with F3-8 EVs

### Slide 4
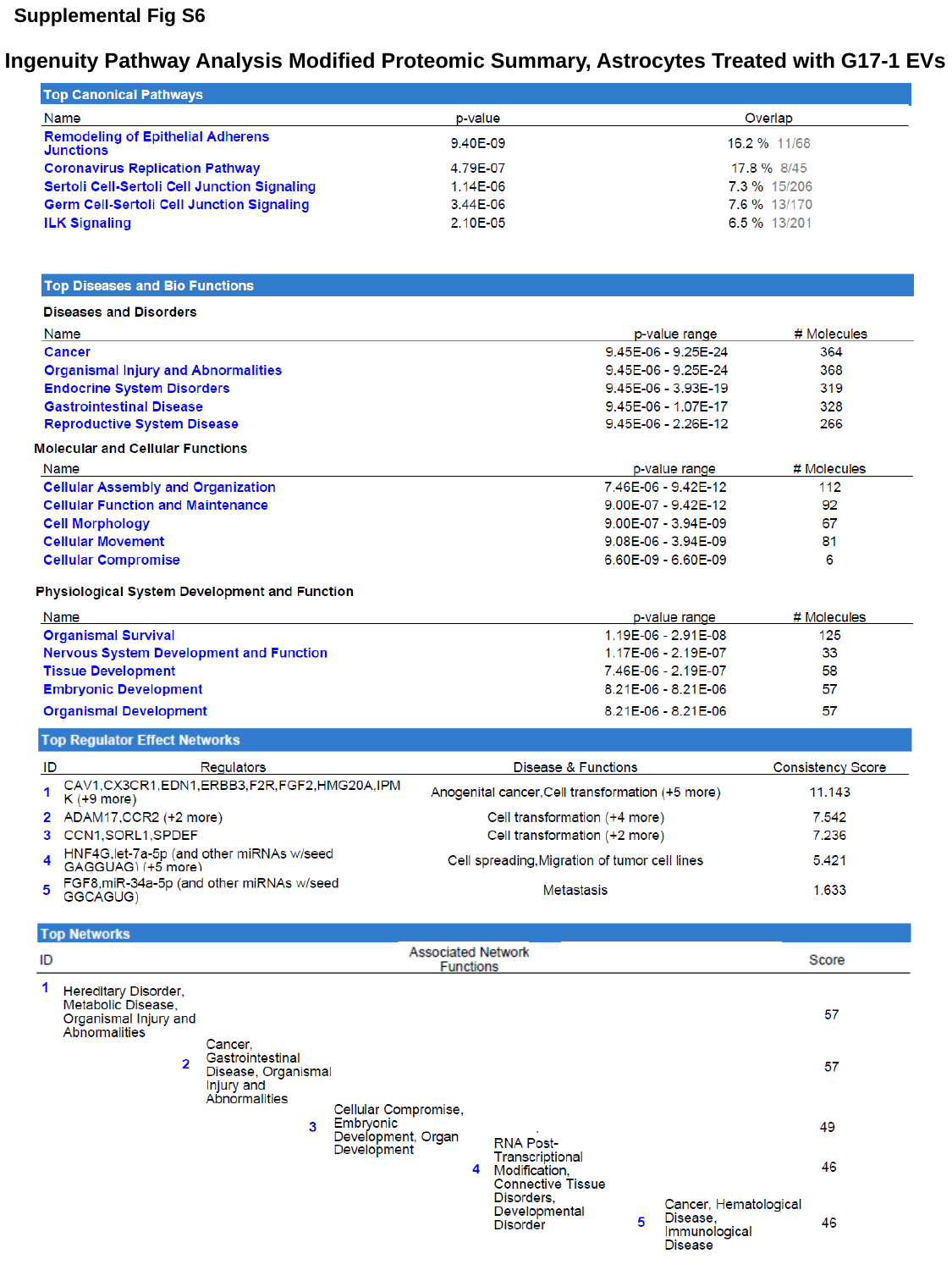

Supplemental Fig S6
Ingenuity Pathway Analysis Modified Proteomic Summary, Astrocytes Treated with G17-1 EVs
