## Supplementary Figure 7 for "A tale of two tumors: differential, but detrimental, effects of glioblastoma extracellular vesicles (EVs) on normal human brain cells"

### Slide 1
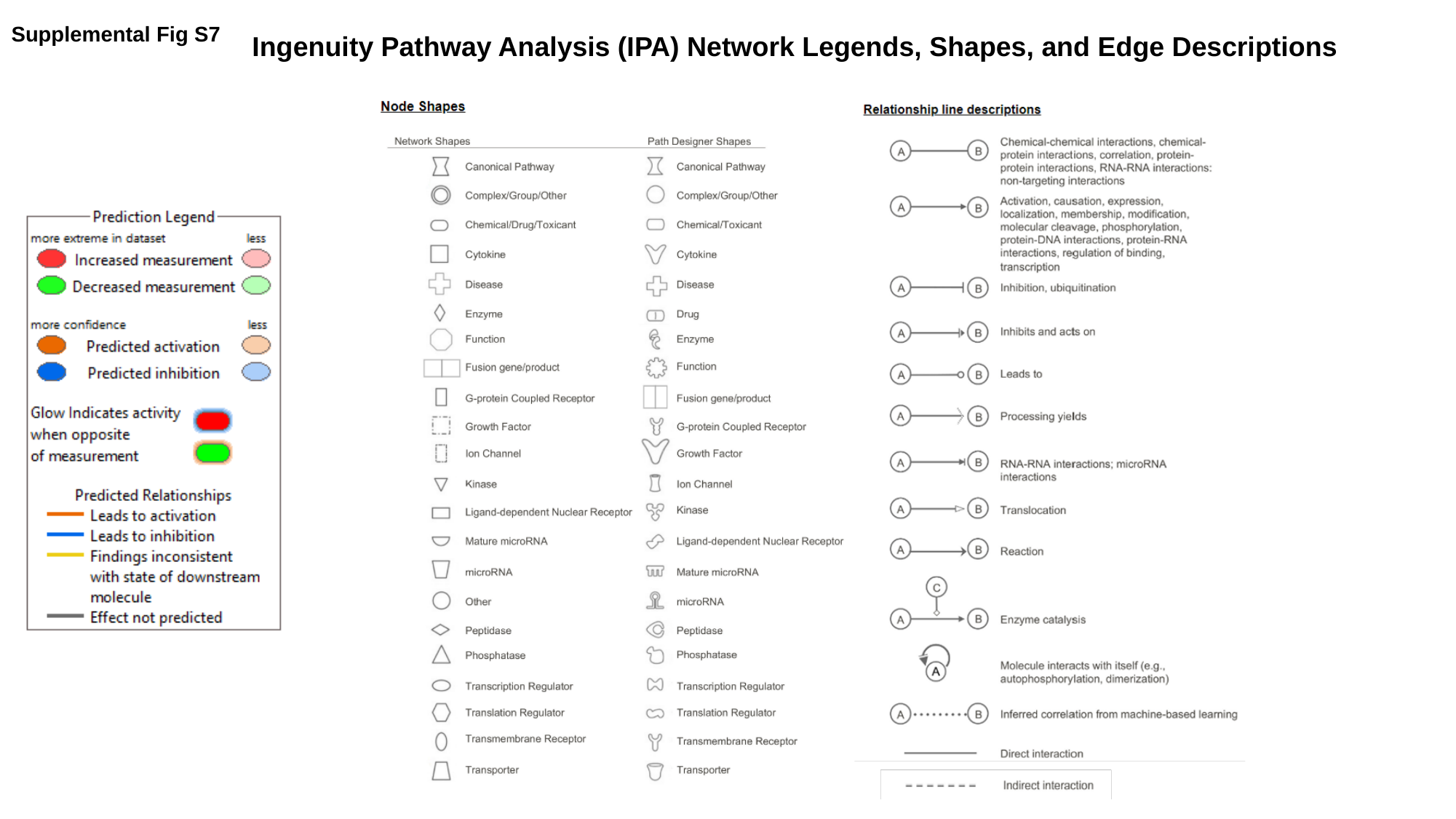

Supplemental Fig S7
Ingenuity Pathway Analysis (IPA) Network Legends, Shapes, and Edge Descriptions
