## Supplementary Figure 8 for "A tale of two tumors: differential, but detrimental, effects of glioblastoma extracellular vesicles (EVs) on normal human brain cells"

**Supplemental Figure S8**

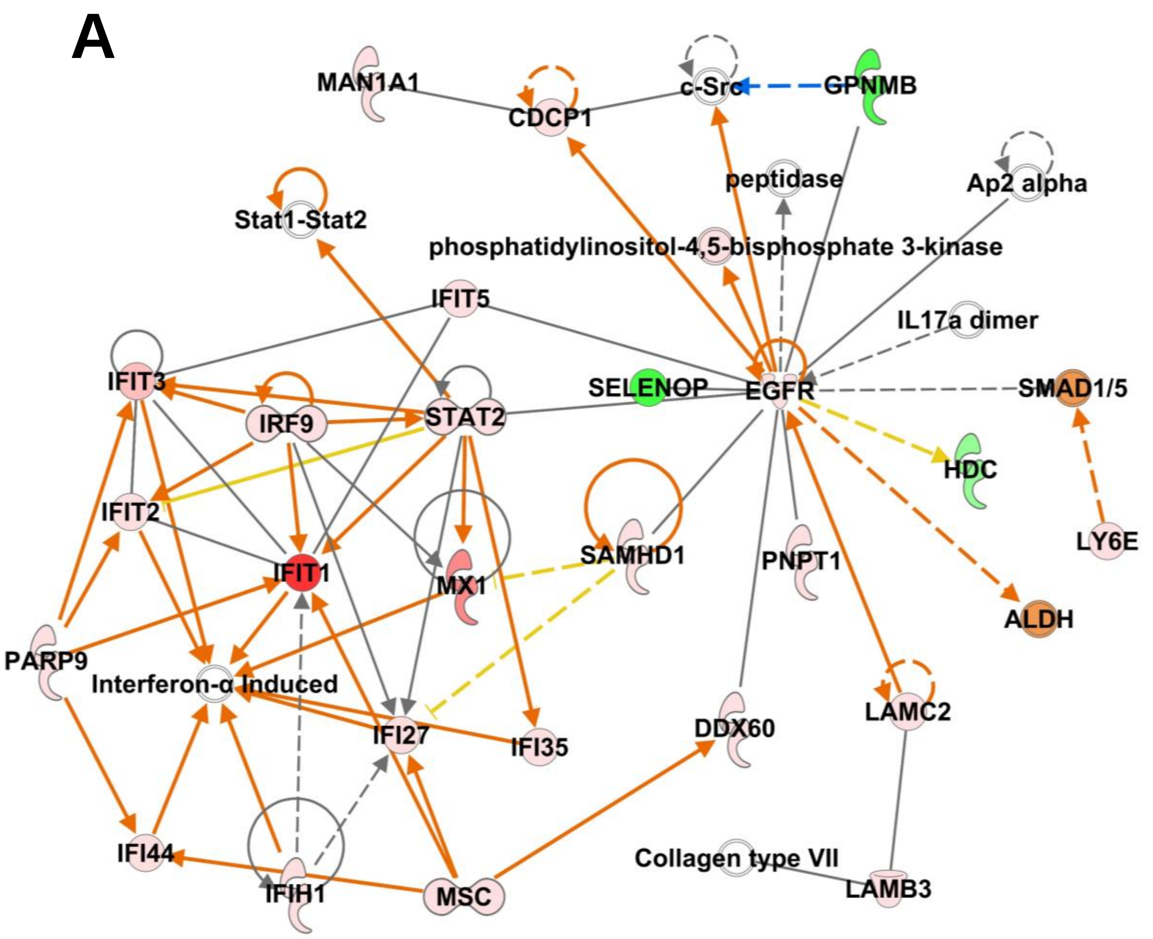


Network 3
Antimicrobial Response, Infectious Diseases,
Inflammatory Response; Score = 38; Focus Molecules = 25

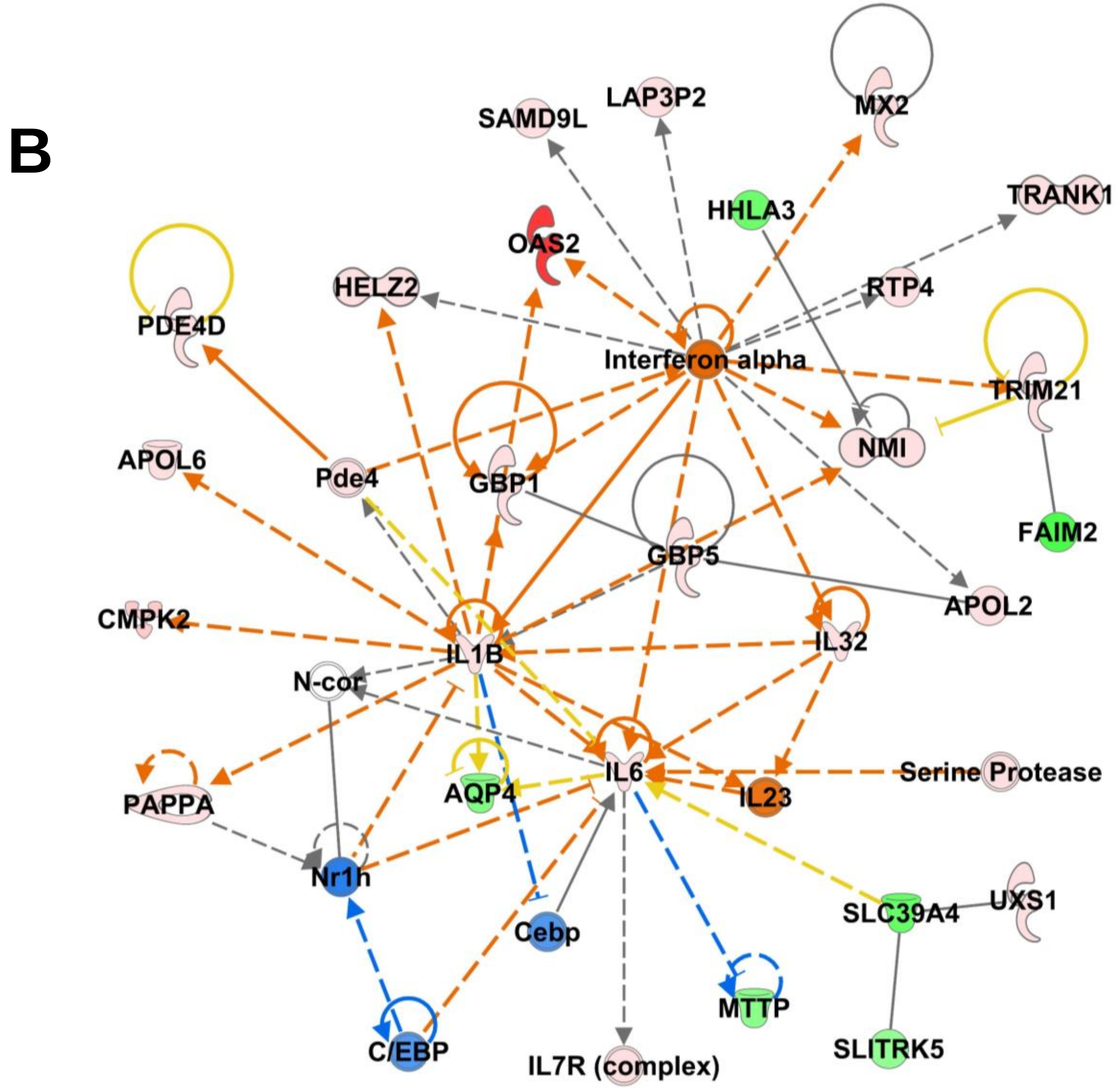


Network 1
Cell Cycle, Cell-to-Cell Signaling and Interaction,
Cellular Development; Score = 41; Focus Molecules = 26
