## Supplementary Figure 9 for "A tale of two tumors: differential, but detrimental, effects of glioblastoma extracellular vesicles (EVs) on normal human brain cells"

### Slide 1
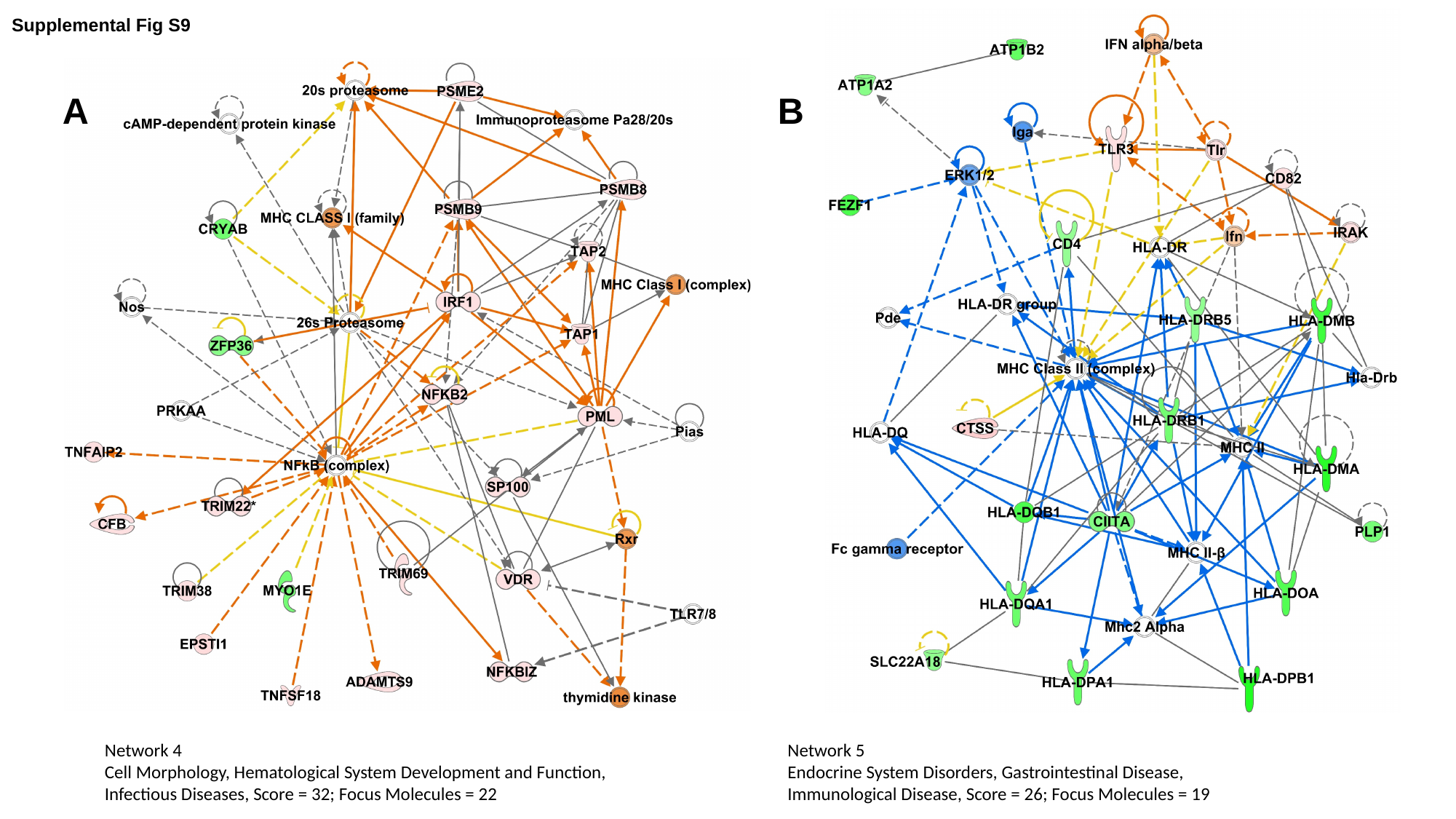

Supplemental Fig S9
A
B
Network 5
Endocrine System Disorders, Gastrointestinal Disease,
Immunological Disease, Score = 26; Focus Molecules = 19
Network 4
Cell Morphology, Hematological System Development and Function,
Infectious Diseases, Score = 32; Focus Molecules = 22
